# Live-cell, single-molecule imaging resolves the dual nature of mRNA silencing by SARS-CoV-2 Nsp1

**DOI:** 10.64898/2026.10.23.759733

**Authors:** Adam Koch, O’Neil Wiggan, Emry Burns, Brian Geiss, Christopher D. Snow, Tatsuya Morisaki, Timothy J. Stasevich

## Abstract

SARS-CoV-2 nonstructural protein 1 (Nsp1) is a conserved viral factor that potently silences host transcripts by binding ribosomes. Yet how it acts at the level of individual mRNAs remains unclear, with competing models arguing that Nsp1 blocks translation initiation, stalls ribosome elongation, cleaves mRNA, or does all three. Here, we use Nascent Chain Tracking to quantify in living cells how Nsp1 reshapes mRNA abundance and translation dynamics with single-molecule resolution. Using reporter transcripts that vary in UTR architecture, initiation mode, and open reading frame length, we find that Nsp1 mainly silences our reporter transcripts by reducing their numbers rather than inhibiting initiation. Two routes to Nsp1-mediated mRNA loss are identified, both requiring cap-dependent initiation: an early route acting during or shortly after initiation and a later route associated with downstream translation that depends in part on ZNF598. Together, these two mechanisms make Nsp1 an efficient and robust suppressor of host gene expression. We further find that reporters containing the SARS-CoV-2 UTRs remain susceptible to Nsp1-mediated mRNA loss, but the 5’ and 3’ UTRs cooperate to increase translation output. Neither viral UTR alone increased translation beyond the host reporter, whereas combining both UTRs substantially enhanced translation in the absence of Nsp1. Together, our results reveal that Nsp1 uses dual, cap-dependent routes to promote mRNA loss across the translation cycle, while the SARS-CoV-2 UTRs provide a cooperative translational advantage rather than robust protection from Nsp1.

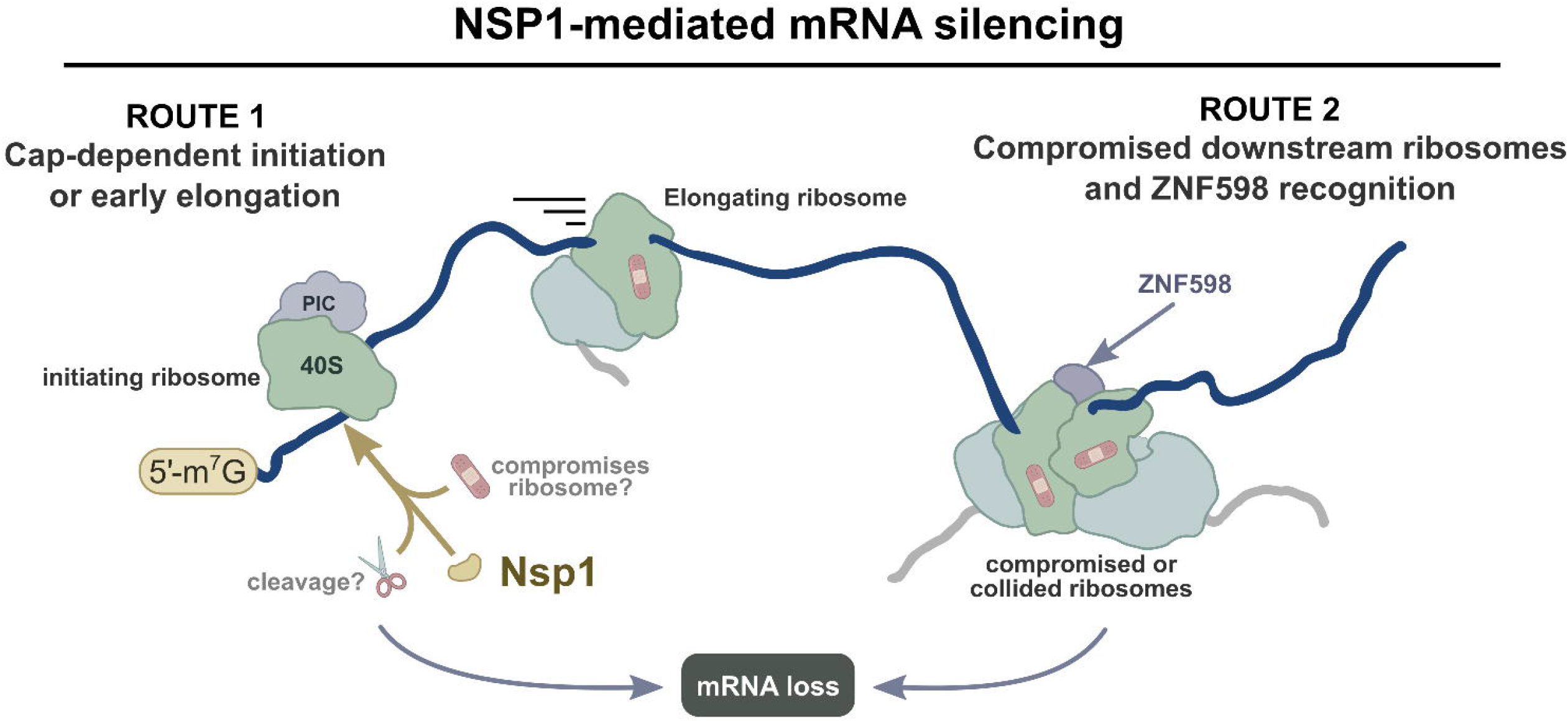

## INTRODUCTION

SARS-CoV-2, the causative agent of the COVID-19 pandemic, has caused a sustained global health burden and widespread societal and economic disruption^1^, with >7 million deaths reported to WHO^2^ and ongoing morbidity that continues to strain health systems^3^. These impacts underscore the need to understand viral pathways that disable host defenses, both to expand therapeutic options for COVID-19 and to strengthen preparedness against future zoonotic coronaviruses. As a positive-sense single-stranded RNA virus, the SARS-CoV-2 genome can be translated directly upon entry into the cytoplasm. Translation begins with open reading frame1a/1ab, placing nonstructural protein 1 (Nsp1) at the N terminus of the first viral polyprotein and making it one of the earliest viral proteins produced. Nsp1 is a conserved coronavirus factor that suppresses host gene expression by silencing mRNAs^4–8^. Despite extensive biochemical and structural work^9,10^, how Nsp1 achieves this at the level of individual transcripts has not yet been resolved.

Part of the difficulty comes from the multiple ways Nsp1 is thought to suppress transcripts. According to structural and reconstitution studies, the C-terminal domain of Nsp1 binds the human 40S ribosomal subunit and occludes the mRNA entry channel, blocking translation initiation^5,9^. In addition, recent *in vitro* work suggests Nsp1 associates with ribosomes to induce mRNA cleavage near the 5′ end^11,12^. No consensus has emerged on whether these activities act independently or in concert, nor on where in the translation cycle they take hold. Some studies report that mRNA loss requires the transcript to be engaged by ribosomes, and that blocking elongation does not prevent it, suggesting Nsp1 may act on elongation-independent pathways or on pathways that may recognize stalled or collided ribosomes^13^. Others find no sign of collisions or translation dependence and report that ribosome binding alone suffices^14^. Precisely when Nsp1 acts on ribosomes is therefore an open question.

Another unresolved question is whether viral transcripts can escape from Nsp1 silencing. Escape has been attributed to the SARS-CoV-2 5′ untranslated region (UTR), with stem-loop 1 in the shared 75-nucleotide leader sequence thought to be necessary and sufficient for evasion^15,16^. Other work, however, finds little evidence for escape. Reporters carrying the viral 5′ UTR are repressed alongside host reporters in cell extracts^5^, and reporters carrying both the viral 5′ and 3′ UTRs are inhibited to the same degree as control human reporters, differing mainly in their higher intrinsic translation efficiency^10^. Complicating matters further, the SARS-CoV-2 3′ UTR is reused across genomic and subgenomic viral RNAs, just as the 5’ leader sequence is^17,18^ It therefore remains unknown if each UTR helps protect viral transcripts from Nsp1 and, if so, whether or not they work together or independently.

Here, to help address these questions, we develop a panel of single-molecule reporters that allow us to track individual mRNA transcripts in living cells and monitor their translation dynamics in the presence and absence of Nsp1. Using this approach, we show that Nsp1 silences reporter transcripts mainly by reducing their numbers rather than lowering their initiation rates. This loss requires cap-dependent initiation and can occur at two stages of translation: first during or shortly after initiation itself, and later in a manner dependent on ZNF598, a well known sensor of ribosome collisions. Consistent with this robust silencing, reporter mRNAs carrying the SARS-CoV-2 UTRs are also reduced by Nsp1. Rather than shielding transcripts from loss, the two UTRs appear to act together to raise the translation of those that remain. More broadly, our single-molecule imaging strategy resolves how a viral shutoff factor acts on individual transcripts, revealing a two-stage mechanism that can help reconcile previously conflicting accounts of Nsp1 function.

## RESULTS

### A system to image and quantify how Nsp1 silences individual mRNAs in living cells

To better resolve how Nsp1 silences host transcripts, we developed a system to track the translation of individual mRNAs in live cells with and without Nsp1. To do this, we used a Nascent Chain Tracking^19^ reporter that contains the 5’ and 3’ UTRs from beta-actin (ActB). This reporter is constitutively expressed and serves as a representative host mRNA (“Host Reporter”). The open reading frame of the reporter contains a tag with ten HA epitopes^20^ followed by luciferase, while the 3’ UTR contains 24 MS2 stem loops^21^. When this reporter is transfected into cells that co-express fluorescent MS2 coat proteins (MCP) to label the mRNA stem loops (HaloTag-MCP) and fluorescent single-chain variable fragments to label the HA epitopes (scFv-mStayGold)^20,22^, individual reporter translation sites appear as distinct puncta where both fluorescent signals are amplified (**Fig. 1A**).

**Figure 1.**
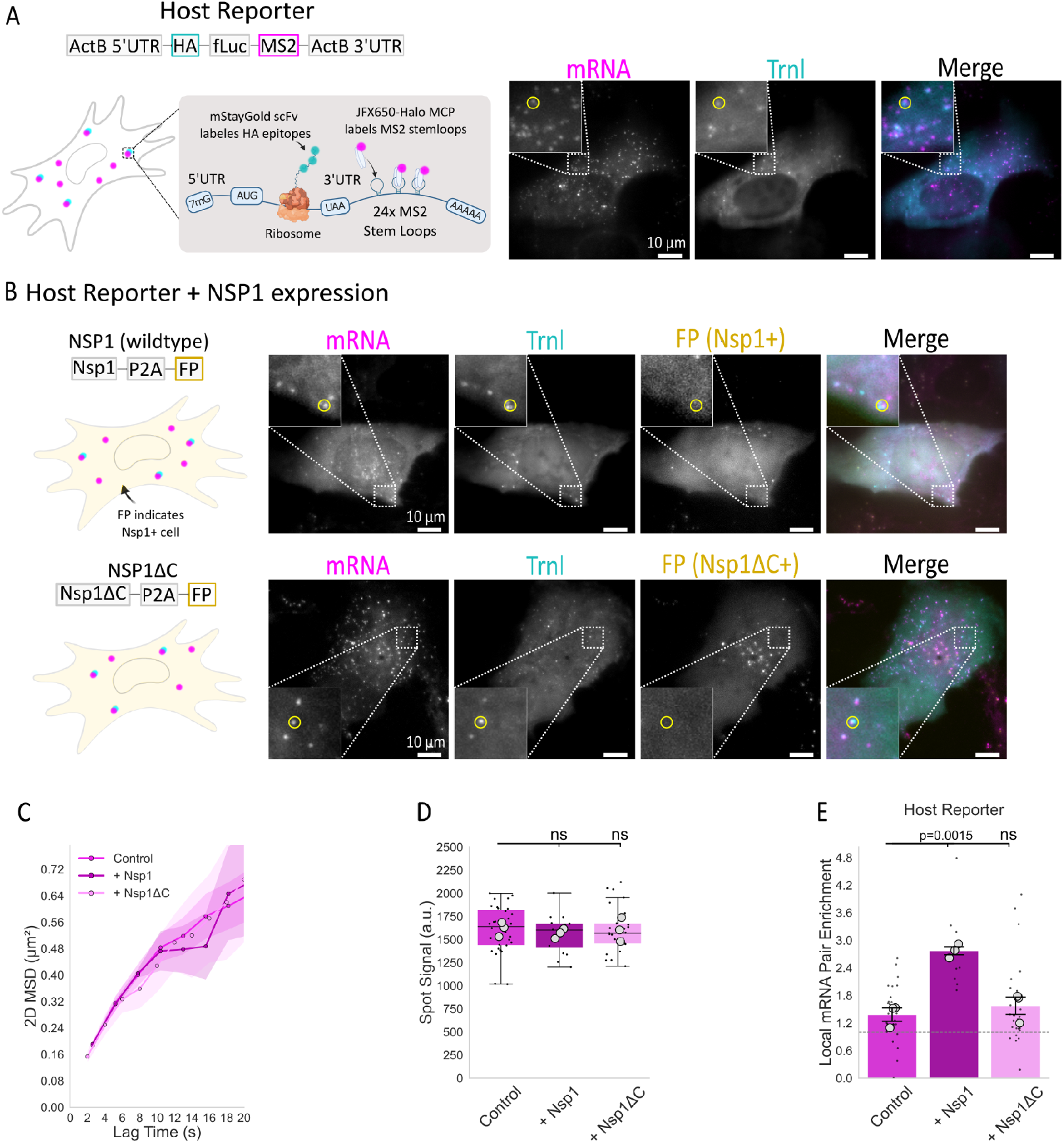
Live-cell imaging strategy for Nsp1 effects on single-mRNA translation. (A) Schematic of the Host Reporter used for Nascent Chain Tracking (NCT). Reporter mRNAs are detected through 24×MS2 stem loops with HaloTag-MCP labeled with JFX650 (magenta), while nascent polypeptides are detected through HA epitopes with anti-HA scFv-mStayGold (cyan). Representative images show individual reporter mRNAs and associated translation sites. Boxed areas are enlarged in the insets, with representative translation sites highlighted (yellow circles). (B) Nsp1 co-expression strategy and representative images of Host Reporter mRNAs in cells expressing WT Nsp1 (top) or the C-terminal deletion mutant Nsp1ΔC (bottom). A P2A-linked fluorescent protein (FP; gold) is uniformly expressed in cells expressing each Nsp1 construct. (C) Mean-squared displacement of reporter mRNAs in control, Nsp1-, and Nsp1ΔC-expressing cells. (D) Distribution of reporter mRNA spot intensities across conditions. (E) Local mRNA pair enrichment relative to spatial randomization. Pair enrichment was 1.4 ± 0.1 (mean ± SEM) in control cells, 2.8 ± 0.1 with Nsp1, and 1.6 ± 0.2 with Nsp1ΔC. Scale bars, 10 μm. P values were calculated using two-sided one-sample t-test for normalized conditions, or paired two-sided t-test for matched conditions across biological replicates. ns indicates p>0.05.

To co-express Nsp1 in cells, we also developed a delivery plasmid. As Nsp1 contains active N and C-terminal domains that a fluorescent tag might disrupt^23,24^, the plasmid encodes untagged wildtype Nsp1 (WT Nsp1) followed by a ribosome skip site (porcine teschovirus 2A; P2A)25 and a separate fluorescent protein (mCherry) that marks Nsp1 positive cells. mCherry fluorescence was used only as a binary marker to identify cells expressing the delivery construct. As a negative control, we also created a similar delivery plasmid that expresses a C-terminal deletion mutant of Nsp1 that cannot bind the ribosome (Nsp1ΔC)^24,26^.

With these plasmids in hand, we could image individual mRNA translation sites in cells under three conditions: (1) without Nsp1 co-expression, (2) with WT Nsp1 co-expression, and (3) with Nsp1ΔC co-expression. Consistent with previous work using luciferase-based reporters^10^, WT Nsp1 coexpression did silence the expression of our Host Reporter at 24 hours (**Fig. S1E**). However, as translation sites were harder to find at this relatively late time point and mRNA tended to cluster (**Movie S1**), we chose to focus our single molecule imaging at an earlier time point, approximately 12 hours after Nsp1 co-expression, when silencing was not yet complete. At this earlier time, we could detect individual mRNA puncta with translation signals in all conditions (**Fig. 1A,B** and **Movie S2**). We verified these puncta were bonafide translation sites by adding the translational inhibitor puromycin, which splits ribosomes^27^. This resulted in a rapid loss of the translation signal within seconds (**Fig. S1A-C**). We also verified that Nsp1 does not significantly change the mobility of mRNAs at this time point (**Fig. 1C** and **S1C**) nor their propensity to aggregate (**Fig. 1D**), although we did measure an increase in the frequency of mRNA pairs within a defined distance compared to random (**Fig. 1E**), suggesting a minor redistribution of mRNAs within the cell (and consistent with the clusters we observed at 24 hours). Together, this work establishes a system with the single-molecule sensitivity needed to more precisely resolve how Nsp1 silences host mRNAs in living cells.

### Nsp1 impacts reporter mRNAs mainly by reducing their number

With our imaging system validated, we first asked how Nsp1 impacts individual host-like mRNAs at the 12 hour time point. Previous studies have shown that Nsp1 can suppress gene expression both by inhibiting translation initiation through its interaction with the ribosome^9^ and by promoting mRNA degradation or cleavage^11,12^. However, the relative contribution of these effects to silencing individual transcripts in cells remains unclear. We therefore compared cells without Nsp1, with WT Nsp1, and with Nsp1ΔC, quantifying Host Reporter mRNA abundance, the fraction of mRNAs undergoing translation, and the translation signal per mRNA. The largest effect was on mRNA abundance: WT Nsp1 reduced the average number of detectable mRNAs per cell by ∼45%, whereas Nsp1ΔC had no significant effect (**Fig. 2A,B**). Translation was affected more modestly. The fraction of mRNAs being translated increased slightly with WT Nsp1, but a similar increase was seen with Nsp1ΔC (**Fig. 2C**), indicating that this effect does not require the C-terminal ribosome-binding domain. Individual translation sites were also ∼60% brighter with WT Nsp1 but unchanged with Nsp1ΔC (**Fig. 2D**). According to our calibration (**Fig. S2**), this corresponds to an increase in ribosome occupancy from ∼9 ribosomes per mRNA to ∼14.

**Figure 2.**
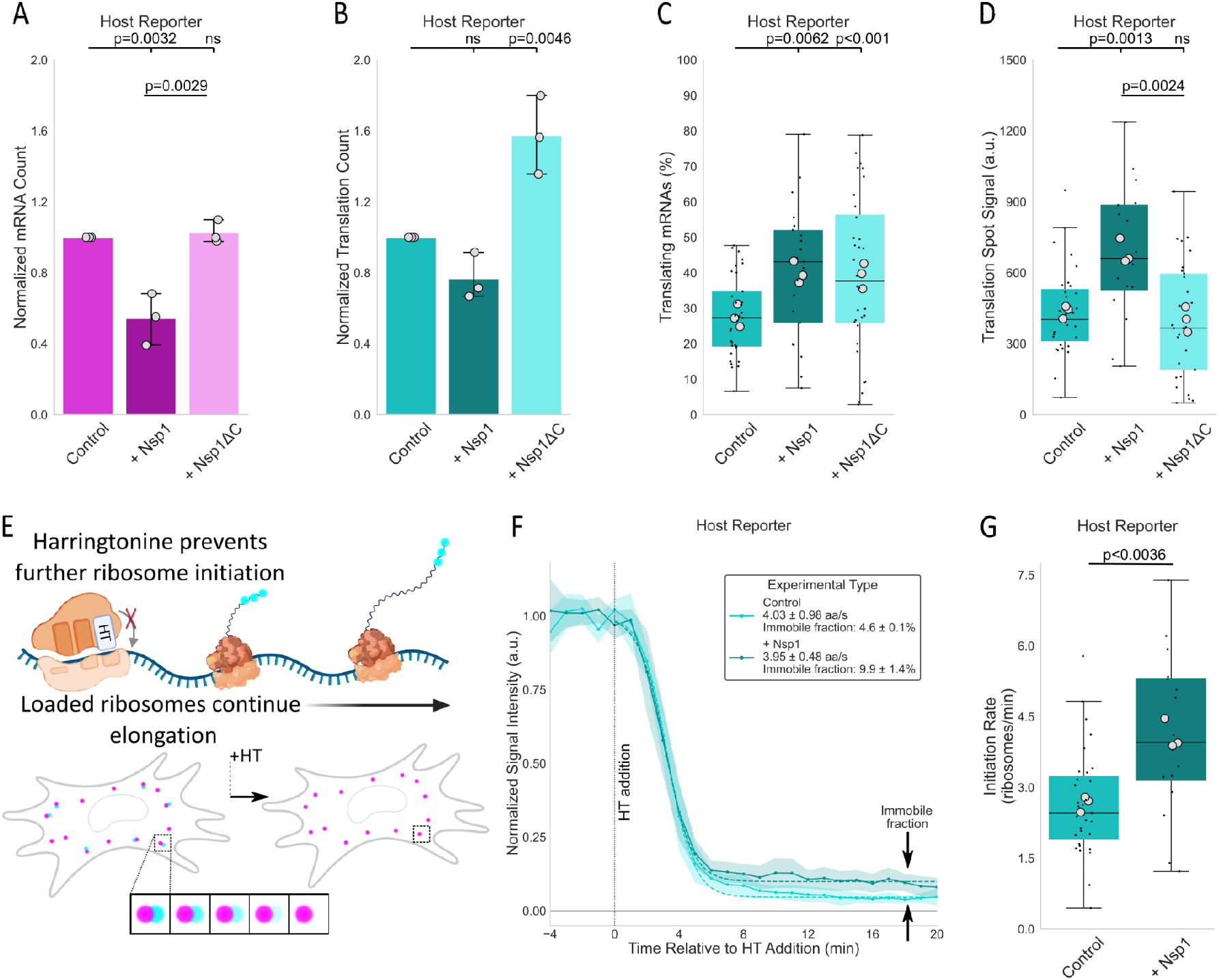
Nsp1 reduces Host Reporter mRNA abundance while increasing initiation rate on surviving mRNAs. (A) Normalized reporter mRNA abundance per cell. Reporter mRNA abundance was 54.2 ± 8.4% (mean ± SEM) of control with WT Nsp1. (B) Normalized number of translation sites per cell. Translation-site number was 76.6 ± 7.6% of control with WT Nsp1. (C) Fraction of reporter mRNAs associated with a detectable translation site. Translating fractions were 39.9 ± 1.8% and 39.2 ± 2.1% for Nsp1 and Nsp1ΔC, respectively. (D) Translation-site intensity of actively translating reporter mRNAs. Translation-site intensity was 437 ± 16 a.u. in control cells and 686 ± 31 a.u. with WT Nsp1. (E) Schematic of the harringtonine (HT) ribosome runoff assay. HT traps newly initiating ribosomes at the start codon, preventing further initiation while allowing already elongating ribosomes to complete translation. (F) Mean normalized translation-site intensity following HT addition. Elongation rates were 4.0 ± 1.0 and 4.0 ± 0.5 aa s⁻¹ for control and Nsp1, respectively. HT-resistant fractions were 4.6 ± 0.1% and 9.9 ± 1.4%. Shading indicates SEM. (G) Estimated ribosome initiation rates calculated from translation-site intensity and nascent-chain calibration. Initiation rates were 2.66 ± 0.7 and 4.1 ± 0.6 ribosomes min⁻¹ for control and Nsp1, respectively. P values were calculated using two-sided one-sample t-test for normalized conditions, or paired two-sided t-test for matched conditions across biological replicates. ns indicates p>0.05.

A larger number of ribosomes per mRNA can arise in two ways: either faster initiation, which would load more ribosomes onto each mRNA, or slowed elongation, which would cause ribosomes to pile up behind one another. To distinguish these possibilities, we measured elongation rates directly using a ribosome runoff assay. We treated cells with Harringtonine, which blocks new rounds of initiation^28^. Elongating ribosomes are unaffected and continue translating until they reach the stop codon and run off the transcript^29,30^. As they do, the nascent chain signal gradually decreases over time (**Fig. 2E** and **Movie S3**). The rate of this decrease can then be fit to estimate the ribosome elongation speed. According to our fits (**Fig. 2F**), Nsp1 co-expression did not have a significant effect on translation elongation speed, which remained ∼4 aa/sec irrespective of Nsp1 presence, although we did measure a small increase in the immobile fraction (**Fig. 2F**). Since there was no major impact on elongation by Nsp1, we conclude that the ribosome initiation rate instead must have increased. Combining the calibrated number of ribosomes per transcript with the fitted elongation rate and the total length of the ORF, we calculate the ribosome initiation rate (see Methods) increased from 2.66 ± 0.68 ribosomes/min to 4.12 ± 0.62 ribosomes/min (**Fig. 2G**). These data suggest, at least at the 12 hour time point, the translation initiation rate increases for Host Reporter in the presence of Nsp1, perhaps reflecting the growing pool of free translation machinery as endogenous genes are broadly silenced.

Together, these experiments suggest that Nsp1 mainly acts by reducing the number of Host Reporter mRNAs per cell rather than reducing their ribosome elongation or initiation rates. These data contradict the model in which Nsp1 directly inhibits translation initiation, which would have depleted the pool of translating mRNAs and also dimmed translation sites.

### Cap-dependent initiation is required for Nsp1-mediated mRNA loss

Because WT Nsp1 reduced Host Reporter mRNA levels while increasing ribosome initiation rates on those that remained, we asked whether translation itself might be the driver of mRNA loss. To test this, we created a single-mRNA reporter that ribosomes cannot translate. Recent work from Shehata et al. showed that Nsp1-dependent decay requires a ribosome engagement with mRNA^13^. According to their data, a transcript that cannot engage ribosomes should be protected from Nsp1. With this in mind, we removed the entire open reading frame from the Host Reporter and replaced the 5’UTR with twelve BoxB stem loops to block ribosome scanning and initiation^31^ (“No ORF Reporter”). Consistent with Shehata et al., co-expressing this reporter with WT Nsp1 did not reduce its mRNA levels **(Figs. 3A**). Thus, preventing ribosome engagement protects the reporter from Nsp1-mediated loss.

**Figure 3.**
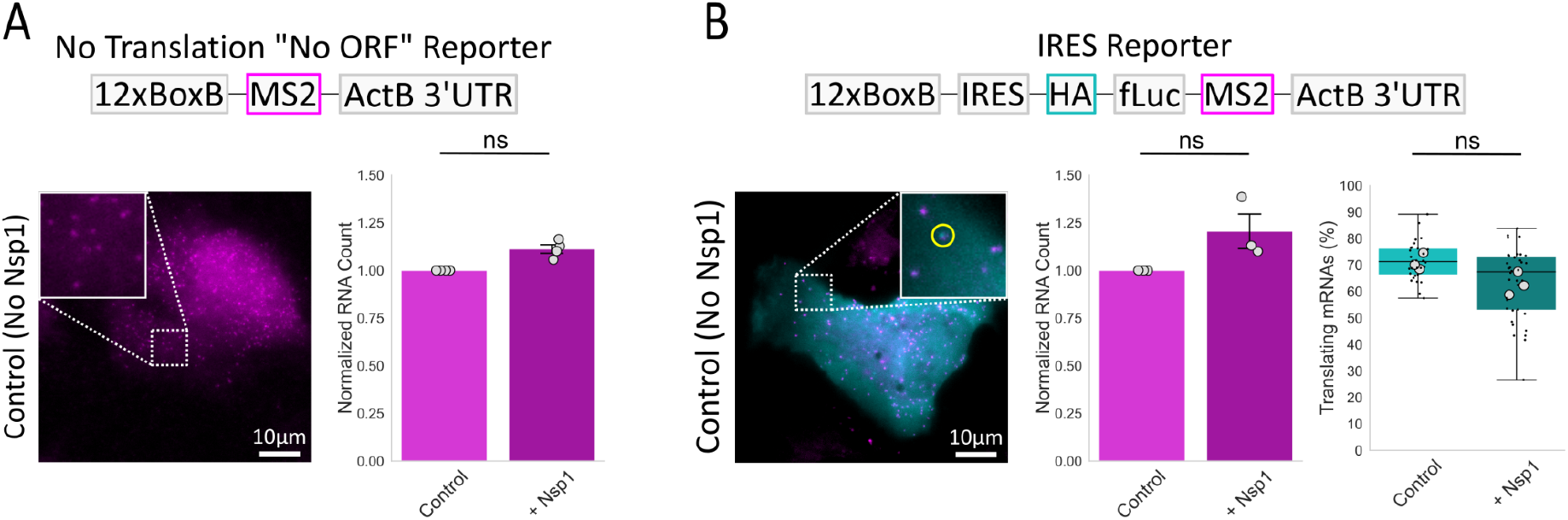
Cap-dependent initiation is required for Nsp1-mediated mRNA loss. (A) Schematic (top), representative image (lower left), and normalized mRNA abundance (lower right) of the No ORF Reporter. The coding region was removed and 12×BoxB stem loops replaced the 5′ UTR to prevent productive ribosome engagement. Representative image shows individual reporter mRNAs. Boxed areas are enlarged in the insets. Normalized reporter abundance with Nsp1 was 111 ± 2.3% (mean ± SEM) of control. (B) Schematic (top), representative image (lower left), normalized mRNA abundance (lower middle), and translating fraction (lower right) of the IRES Reporter. Cap-dependent scanning was blocked with 12×BoxB stem loops, while the EMCV IRES drove cap-independent translation of the downstream ORF. Representative images show individual reporter mRNAs and associated translation sites. Boxed areas are enlarged in the insets, with representative translation site highlighted (yellow circle).Normalized reporter abundance with Nsp1 was 120 ± 9.1% of control. Translating fractions were 71 ± 1.8% in control cells and 62.8 ± 2.5% with Nsp1. Scale bars, 10 μm. P values were calculated using two-sided one-sample t-test for normalized conditions, or paired two-sided t-test for matched conditions across biological replicates. ns indicates p>0.05.

To determine which step of translation renders an mRNA susceptible to Nsp1, we next asked if cap-dependent initiation is necessary to cause Nsp1 sensitivity. In vitro studies have shown that ribosome-bound Nsp1 can promote endonucleolytic cleavage near the 5’ end of mRNAs during early ribosome engagement^11,12^, coupling cleavage to cap-dependent initiation. If cap-dependent initiation is required for Nsp1 susceptibility, then a transcript translated in a cap-independent manner should remain protected, regardless of translation state. To test this, we used the internal ribosome entry site (IRES) from the encephalomyocarditis virus (EMCV) to drive the translation of the Host Reporter ORF in a cap-independent way^32,33^. We again added BoxB stem loops in the 5′ UTR to block any residual cap-dependent initiation, so that translation could only proceed through the IRES (“IRES Reporter”). Approximately 70% of the reporter mRNAs were actively translating, but the low translation-site signal indicated each mRNA is translated by relatively few ribosomes (**Fig. S3C)**. This fraction was unchanged by Nsp1 (**Fig. 3B, right** and **Fig. S3A-D**). Yet despite this persistent translation, Nsp1 did not reduce the reporter mRNA abundance (**Fig. 3B, middle**). These data therefore demonstrate that cap-dependent initiation is the critical interaction required for the loss of our reporter by Nsp1.

### Nsp1-mediated mRNA loss requires cap-dependent initiation but also scales with ORF length

Having shown cap-dependent initiation is required for Nsp1-mediated mRNA loss, we next asked how quickly Nsp1 can act upon initiation. To resolve this, we modified our No ORF reporter by inserting upstream open reading frames (uORFs; 24–80 aa in length from the *EMI1* 5′ UTR^34^) in front of the BoxB stem loops, restricting translation to these short uORFs alone. With this “uORF Only Reporter,” ribosomes could only briefly translate before terminating, giving Nsp1 only a short window after initiation to act. When this reporter was co-expressed with WT Nsp1, mRNA abundance dropped by ∼35% (**Fig. 4A**), indicating that Nsp1 can act within this brief window of time. As a control, we generated a matched construct in which the start codons of all uORFs were mutated to prevent initiation altogether (NoStart uORF Reporter). This construct showed no reduction in mRNA levels (**Fig. 4B**), confirming that initiation at the start codon is required to open the window for Nsp1 action.

**Figure 4.**
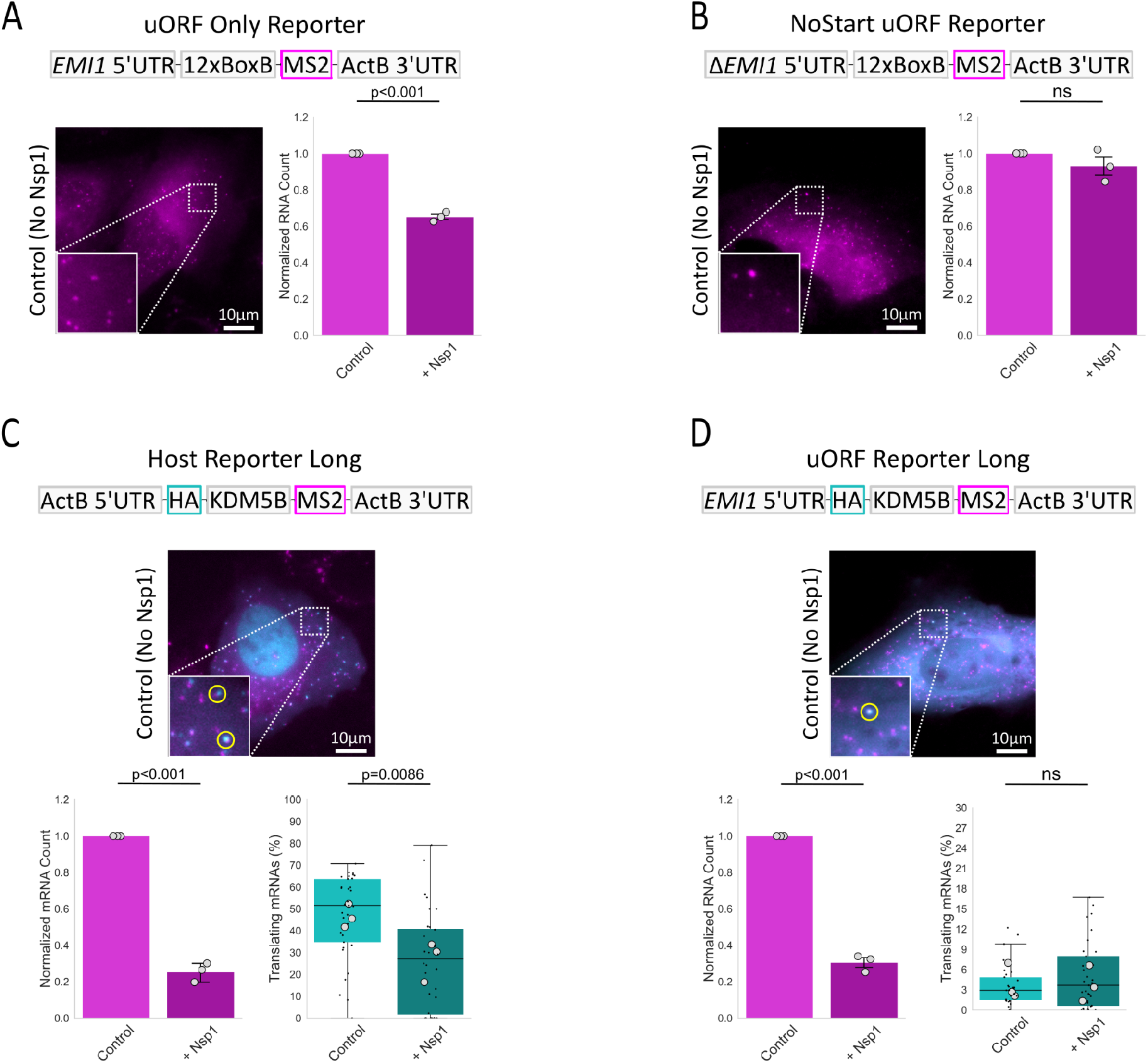
Downstream translation increases Nsp1-mediated mRNA loss. (A) Schematic (top), representative image (lower left), and normalized mRNA abundance (lower right) of the uORF Only Reporter. The *EMI1* 5′ UTR permits initiation within upstream open reading frames, while downstream BoxB stem loops prevent productive translation of a main ORF. Normalized reporter abundance with Nsp1 was 65.2 ± 1.5% (mean ± SEM) of control. Representative images show individual reporter mRNAs. Boxed areas are enlarged in the insets. (B) Same as (A), but for the NoStart uORF Reporter, in which uORF start codons were mutated to prevent initiation. Normalized mRNA abundance with Nsp1 was 93.1 ± 5.1% of control. (C) Schematic (top) of Host Reporter Long containing the KDM5B coding region. Representative images (middle) show individual reporter mRNAs and associated translation sites. Boxed areas are enlarged in the insets, with representative translation sites highlighted (yellow circle). Bottom, normalized mRNA abundance with Nsp1 was 25.4 ± 3.0% of control. Translating fractions were 46.5 ± 3.1% in control cells and 26.8 ± 5.3% with Nsp1. (D) Same as (c), but for the uORF Reporter Long containing the KDM5B coding region. Normalized mRNA abundance with Nsp1 was 30.5 ± 2.7% of control, with translating fractions of 3.9 ± 1.5% and 3.8 ± 1.5% for control and Nsp1, respectively. Scale bars, 10 μm.P values were calculated using two-sided one-sample t-test for normalized conditions, or paired two-sided t-test for matched conditions across biological replicates. ns indicates p>0.05.

Although these experiments show that Nsp1 can act quickly, the mRNA loss triggered by the uORF Only Reporter (∼35%) was less than what we observed for the Host Reporter (∼45%). This left us wondering if the difference reflects Nsp1-triggered events associated with the translation of the main ORF, rather than with earlier events linked to initiation itself. If this were the case, we would predict that transcripts with longer ORFs (which take longer to translate and therefore give Nsp1 more time to act) would generally be hit harder by Nsp1 than those with shorter ORFs. To directly test this idea, we created longer versions of both the Host Reporter and the uORF Only Reporter. Specifically, we replaced the 550-aa firefly luciferase ORF in the Host Reporter with the much longer KDM5B ORF (1550 aa), creating “Host Reporter Long” (**Fig. 4C** and **S4A-D**). Similarly, we replaced the BoxB stem loops in the uORF Only Reporter with the KDM5B ORF, creating “uORF Reporter Long” (**Fig. 4D** and **S4E-H**). In both cases, the longer main ORF significantly increased Nsp1-mediated mRNA loss, from ∼45% to ∼75% for Host Reporter Long and from ∼35% to ∼70% for uORF Reporter Long, confirming that longer ORFs are hit harder by Nsp1. Notably, this increase was similar in magnitude for both reporters, even though uORF Reporter Long is translated less frequently and by fewer ribosomes than Host Reporter Long (**Fig. S4C** vs **S4G)**, indicating that high ribosome occupancy is not required for Nsp1-mediated mRNA loss. Together, these results indicate that cap-dependent initiation alone cannot account for all Nsp1-mediated mRNA loss. Instead, the length of the main ORF also contributes, even when that ORF is only rarely translated.

### Nsp1-mediated mRNA loss depends in part on ribosome-associated quality control

Our data so far show that extending the translated region increases Nsp1-mediated mRNA loss, although this loss still requires cap-dependent initiation at a start codon. Two mechanisms could explain this length dependence. First, the increased susceptibility of longer transcripts could reflect an intrinsic property of the transcript itself, independent of ribosome elongation, such as an increase in its flexibility, which could lower its overall stability. Alternatively, ribosomes translating longer ORFs may have more opportunities to enter stalled or otherwise problematic states in the presence of Nsp1. Persistent stalls can engage ribosome-associated quality control (RQC) pathways that promote degradation of the associated mRNA^35^. Although one might expect such stalling-associated loss to scale with ribosome occupancy, this relationship may not be so simple: recent work has shown that neighboring ribosomes can help resolve stalls through transient collisions, a process termed ribosome cooperativity^36^, so that higher ribosome density does not necessarily translate into a higher probability of persistent, problematic collisions.

To distinguish these hypotheses, we repeated experiments in cells lacking ZNF598^37^ (**Fig. S5**), an early RQC factor that is thought to signal problematic mRNAs by binding the unique interface formed by collided ribosomes^38^. If the additional loss associated with longer ORFs arises from ZNF598-sensitive events on elongating ribosomes, knockout of ZNF598 should preferentially protect the longer reporters. Conversely, if the increased susceptibility is an intrinsic property of the longer transcript, knockout of ZNF598 should have little effect on mRNA levels. In ZNF598-knockout cells (**Movies S4** and **S5)**, Nsp1-mediated mRNA loss decreased from ∼45% to ∼21% for the Host Reporter (**Fig. 5A** vs. **2A**) and from ∼75% to ∼5% for the Host Reporter Long (**Fig. 5B** vs. **4C**).

**Figure 5.**
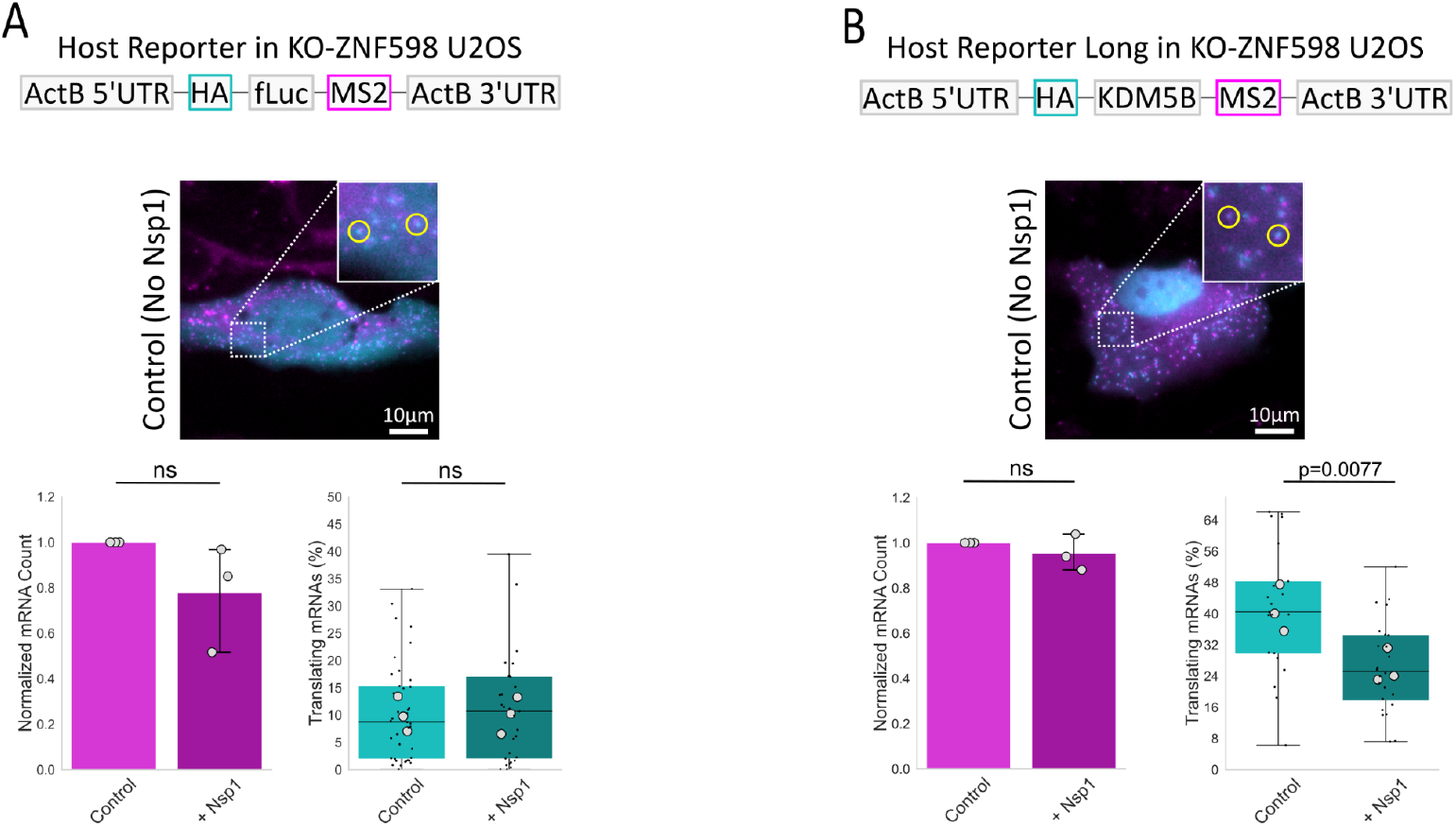
ZNF598 knockout rescues Nsp1-mediated mRNA loss. (A) Host Reporter measurements in ZNF598-knockout U2OS cells. Top, schematic of Host Reporter. Middle, representative image shows individual reporter mRNAs and associated translation sites. Boxed area is enlarged in the inset, with representative translation sites highlighted (yellow circle). Bottom, normalized reporter mRNA abundance with Nsp1 was 79.0 ± 1.0% (mean ± SEM) of control. Translating fractions were 10.1 ± 1.9% and 10.0 ± 2.0% for control and Nsp1, respectively. (B) Same as (A), but for Host Reporter Long in ZNF598-knockout cells. Normalized mRNA abundance with Nsp1 was 95.2 ± 4.6% of control. Translating fractions were 41.1 ± 3.5% and 26.1 ± 2.6% for control and Nsp1, respectively. Scale bars, 10 μm. P values were calculated using two-sided one-sample t-test for normalized conditions, or paired two-sided t-test for matched conditions across biological replicates. ns indicates p>0.05.

Together, these data indicate that a significant portion of Nsp1-mediated loss depends on ZNF598, implicating aberrant ribosome states that arise during downstream translation rather than events during or soon after cap-dependent initiation. Given the established role of ZNF598 in recognizing collided ribosomes^38^, these findings are consistent with problematic ribosome collisions contributing to Nsp1-mediated mRNA loss. Importantly, this pathway may remain active even on sparsely translated mRNAs such as uORF Reporter Long. Because ribosome cooperativity^36^ depends on the presence of neighboring ribosomes, the low ribosome occupancy of uORF Reporter Long may limit this buffering effect, potentially allowing stalls to persist long enough to engage ZNF598-dependent quality control despite the reporter’s low average ribosome density.

### Nsp1-mediated mRNA loss extends to an endogenous gene

Taken together, our data support a model in which Nsp1 promotes mRNA loss, first during our soon after cap-dependent initiation and later through a second, ZNF598-sensitive pathway associated with downstream translation elongation. To test whether this model extends beyond transiently transfected reporters, we next examined an endogenous gene. We used a previously developed HeLa cell line in which sequential rounds of CRISPR/Cas9 editing were used to tag the MYH9 locus at multiple positions^39^. MYH9 encodes the heavy chain of non-muscle myosin-2A, a cytoplasmic actin-based motor that drives cytoskeletal contractility. The transcript is constitutively expressed and heavily translated, with the majority of mRNAs engaged by ribosomes at steady state^39^. The edited MYH9 locus contains an N-terminal 6×FLAG tag for detecting MYH9 translation, a C-terminal mClover tag to label mature myosin-2A, and a 24×MS2 stem loop tag in the 3’UTR to label mRNA (**Fig. 6A**). Similar to our transiently transfected reporters, we can image endogenous MYH9 transcripts using HaloTag-MCP (stably co-expressed in the edited cells), and simultaneously assess their translational status by either coexpressing anti-FLAG scFvs or fixing cells and then staining with anti-FLAG antibodies. The latter approach allowed us to measure translation without co-expressing scFvs, avoiding any influence their expression and binding might have on our measurements.

**Figure 6.**
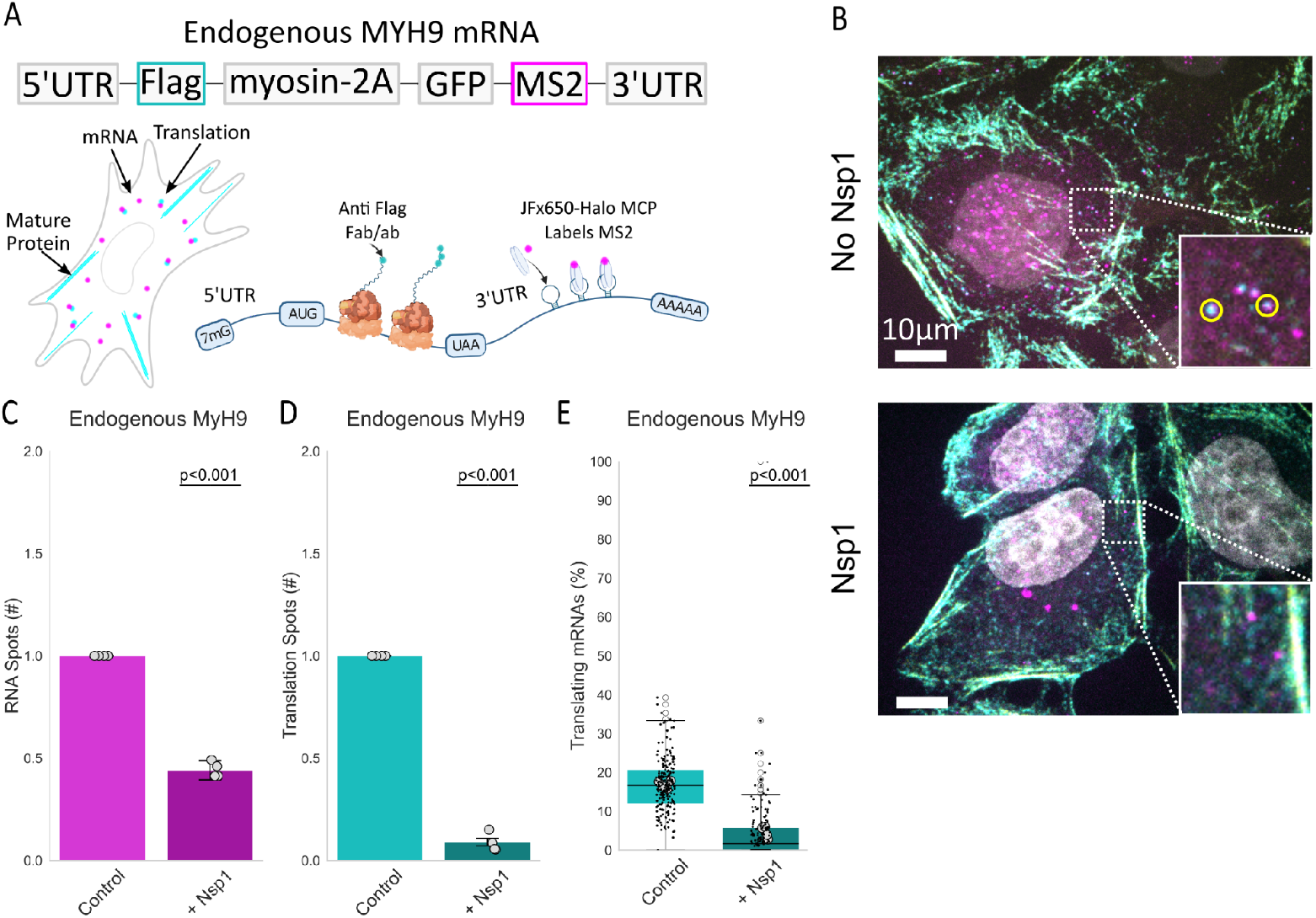
Nsp1 reduces endogenous MYH9 mRNA abundance and translation. (A) Schematic of the endogenously tagged MYH9 locus. MYH9 mRNAs are detected through 24×MS2 stem loops with HaloTag-MCP (magenta), while nascent translation is detected through an N-terminal FLAG tag using either anti-FLAG antibody fragments (Fab) or antibodies (AB; cyan). (B) Representative images of endogenous MYH9 mRNAs and translation sites in control (No Nsp1; top) and Nsp1-expressing cells (bottom). Note that mature myosin-2A is labeled by both anti-FLAG Fab/AB and GFP (channel not shown), in contrast to nascent chains that are only labeled by anti-FLAG Fab (since GFP has not yet matured). Boxed regions are enlarged in insets. (C) Normalized MYH9 mRNA abundance per cell. Normalized mRNA abundance with Nsp1 was 44.4 ± 1.9% (mean ± SEM) of control. (D) Normalized number of MYH9 translation sites per cell. Translation-site number with Nsp1 was 8.7 ± 2.3% of control. (E) Fraction of MYH9 mRNAs associated with detectable translation sites. Translating fractions were 16.9 ± 0.5% in control cells and 4.6 ± 0.8% with Nsp1. Scale bars, 10 μm. P values were calculated using two-sided one-sample t-test for normalized conditions, or paired two-sided t-test for matched conditions across biological replicates. ns indicates p>0.05.

Given the exceptional length of the MYH9 ORF (1960 aa) and its high rate of translation^39^, we predicted it would respond to Nsp1 like Host Reporter Long, which is similar in size (1550 aa) and fraction translating (∼50%; **Fig. 6E**). Consistent with this prediction, WT Nsp1 reduced endogenous MYH9 mRNA abundance by ∼55%, close to the ∼75% we measured with Host Reporter Long (**Fig. 6B,C**). Similarly, the number of translation sites dropped by ∼92% for MYH9, compared to ∼86% (**Fig. 6D,E**) for Host Reporter Long (**Fig. S4B**). These measurements on an endogenous transcript corroborate those from our transiently transfected reporters, and suggest that MYH9, like Host Reporter Long, is targeted by Nsp1 at both the initiation and elongation stages of translation.

### SARS-CoV-2 UTRs cooperate to enhance translation without preventing Nsp1-mediated mRNA loss

Our preceding experiments indicate that Nsp1 can promote transcript loss at two stages of RNA translation, affecting both longer and shorter RNAs on which host ribosomes initiate in a cap-dependent manner. This raises an apparent paradox for SARS-CoV-2 RNAs, which are themselves capped and translated by host ribosomes. Previous studies have reached different conclusions as to how viral transcripts remain expressed under these conditions. Several studies have identified the SARS-CoV-2 5′ leader, and particularly stem-loop 1, as sufficient to promote escape from Nsp1-mediated repression^15,16^, whereas other work found that reporters containing the viral UTRs remain similarly sensitive to Nsp1 and instead differ primarily in their higher intrinsic translation efficiency^10^. We therefore asked whether the SARS-CoV-2 UTRs primarily protect transcripts from Nsp1-mediated loss or instead promote translation through another mechanism. To distinguish these possibilities, we generated three NCT reporters from the Host Reporter backbone: one containing both the full-length SARS-CoV-2 5′ and 3′ UTRs, one containing the SARS-CoV-2 5′ UTR with the ActB 3′ UTR, and one containing the ActB 5′ UTR with the SARS-CoV-2 3′ UTR^40^ (**Fig. 7A**).

**Figure 7.**
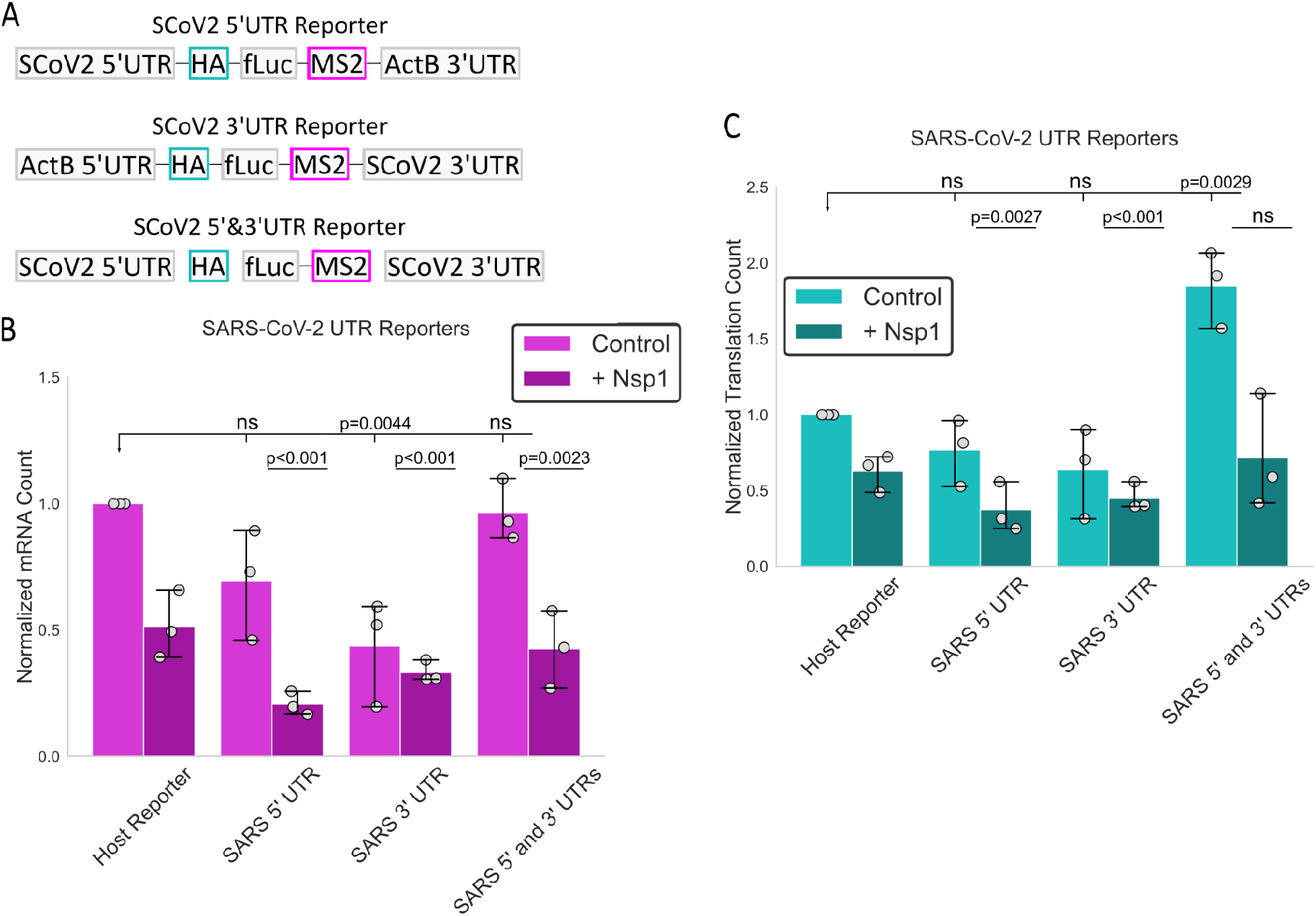
SARS-CoV-2 UTRs differentially modulate Nsp1 sensitivity. (A) Schematics of NCT reporters containing the SARS-CoV-2 5′ UTR, SARS-CoV-2 3′ UTR, or both viral UTRs surrounding the Host Reporter coding region. (B) Normalized reporter mRNA abundance for the Host Reporter and SARS-CoV-2 UTR reporters in control and Nsp1-expressing cells. In Nsp1-expressing cells, normalized mRNA abundance relative to Host Reporter control was 20.7 ± 2.7% (mean ± SEM) for the SARS-CoV-2 5′ UTR reporter, 40.6 ± 11.1% for the SARS-CoV-2 3′ UTR reporter, and 42.5 ± 8.8% for the combined viral UTR reporter. (C) Normalized number of translation sites per cell for each reporter in the absence or presence of Nsp1. In the absence of Nsp1, translation-site abundance for the combined viral UTR reporter was 185 ± 14.7% of control. In Nsp1-expressing cells, normalized translation-site numbers were 37.4 ± 9.4% for the SARS-CoV-2 5′ UTR reporter, 48.4 ± 14.7% for the SARS-CoV-2 3′ UTR reporter, and 71.5 ± 21.8% for the SARS 5′ and 3′ UTRs compared to control. P values were calculated using two-sided one-sample t-test for normalized conditions, or paired two-sided t-test for matched conditions across biological replicates. ns indicates p>0.05.

Transfecting each of these reporters into cells with and without Nsp1 revealed that the viral UTRs did not provide robust protection against Nsp1-mediated mRNA loss. Most notably, the reporter containing both viral UTRs, which most closely approximates natural viral transcripts, decreased by ∼60% in the presence of WT Nsp1, comparable to the Nsp1 sensitivity of the Host Reporter (**Fig. 7B**). The 5′ UTR-only reporter was similarly sensitive, decreasing by ∼70%. The 3’UTR-only reporter was less affected, decreasing by a statistically insignificant ∼24% compared to its control. In the absence of Nsp1, all three viral UTR reporters also had lower mRNA abundance than the Host Reporter, with a significant reduction observed for the 3′ UTR-only reporter. Thus, only the SARS-CoV-2 3′ UTR by itself appears to partially protect our reporter transcripts from Nsp1, although this protection was not maintained when the 5′ and 3′ UTRs were combined.

Although the viral UTRs did not confer resistance to Nsp1-mediated mRNA loss, they cooperated to substantially increase translation output. In the absence of Nsp1, reporters containing either viral UTR individually produced approximately the same number of translation sites per cell as the Host Reporter. In contrast, combining the SARS-CoV-2 5′ and 3′ UTRs nearly doubled the number of translation sites per cell (**Fig. 7C**). Thus, neither viral UTR alone increased translation beyond the host-like reporter, whereas together they produced a substantial translational advantage. Nsp1 reduced translation-site number across all reporter contexts, but the fraction of surviving mRNAs that remained translationally active changed little. The reduction in translation sites therefore largely tracked with mRNA loss rather than complete translational silencing of the remaining transcripts. Together, these results suggest that the principal advantage conferred by the SARS-CoV-2 UTRs in our reporter system is not protection from Nsp1, but their cooperative ability to promote efficient translation. This interpretation is consistent with previous cell-free measurements in which SARS-CoV-2 RNAs containing both viral UTRs showed enhanced intrinsic translation while remaining susceptible to Nsp1^10^.

## DISCUSSION

Nsp1 is widely reported to inhibit host translation, yet the underlying mechanisms have remained difficult to reconcile, with different studies implicating distinct and sometimes apparently contradictory modes of inhibition. Here, we used a panel of modular single-molecule reporters to systematically alter features of individual transcripts and determine how these changes affect their susceptibility to Nsp1. Even in this simplified system, Nsp1 appears to silence mRNAs through multiple routes. One acts during or shortly after translation initiation, consistent with ribosome-dependent cleavage observed in vitro^11,12^, whereas a second emerges during elongation and depends on the ribosome quality-control factor ZNF598^38^.

Unexpectedly, both routes require cap-dependent initiation. This makes sense for the early-acting component of our model as biochemical studies have shown that Nsp1 can promote ribosome-dependent cleavage of mRNAs near their 5′ ends^11,12^. It is less intuitive, however, how the second route, which acts later during elongation, is linked to cap-dependent initiation since elongating ribosomes are physically separated from the 5′ end of the transcript. One possibility is that interaction with Nsp1 during cap-dependent initiation leaves the initiating ribosome in a compromised state that is carried into elongation, increasing its propensity to pause or stall and ultimately trigger ribosome quality control via ZNF598. Consistent with this idea, EMCV IRES-driven translation did not lead to mRNA loss, suggesting that features specific to canonical cap-dependent ribosome recruitment are required for Nsp1-mediated mRNA loss. Although ZNF598 is best characterized as a sensor of collided ribosomes^38^, implying compromised ribosomes collide with one another more often, recent biochemical work indicates that collision is not an absolute requirement for ZNF598-mediated ubiquitination or ASCC-dependent ribosome disassembly^41^. As well, Nsp1 has been reported to associate with ZNF598 and ASCC3, although whether these interactions are direct or mediated through ribosomal complexes remains unclear^42^.

An appealing aspect of a compromised ribosome model is that it may explain why relatively isolated ribosomes can be particularly susceptible to Nsp1-associated defects. Recent single-ribosome imaging showed that neighboring ribosomes can cooperate during translation, with transient ribosome collisions helping resolve pauses and promote processive elongation^36^. In this framework, densely translated mRNAs could paradoxically be less susceptible to elongation-associated Nsp1 effects because neighboring ribosomes help resolve transient pauses before they develop into persistent quality-control-sensitive states. Conversely, isolated ribosomes would lack this cooperative effect, allowing Nsp1-associated elongation defects to persist and become substrates for ZNF598-dependent quality control. Thus, mRNAs with fewer translating ribosomes may actually be more susceptible to elongation-associated mRNA loss by Nsp1.

At first glance, this model appears to differ from previous bulk measurements arguing that Nsp1 does not promote the accumulation of collided ribosomes. Wang et al. found that Nsp1 reduced collision-associated Rps3 ubiquitination and decreased EDF1 and ASCC3 association with polysomes, leading the authors to conclude that Nsp1 promotes collision resolution rather than accumulation^43^. However, these measurements reflect the net behavior of a cellular ribosome pool that is extensively remodeled by Nsp1-mediated translational shutdown^8^. A transient or transcript-restricted stalled state could therefore influence the fate of individual mRNAs without producing a detectable increase in bulk collision markers, especially if problematic mRNAs are quickly removed from cells, as our data would suggest. Moreover, our model does not require the persistent accumulation of large ribosome traffic jams. Instead, it requires only that a subset of translating ribosomes enter a ZNF598-sensitive state long enough to influence mRNA fate. Single-molecule measurements may therefore reveal heterogeneous or short-lived ribosome states that are obscured in population-averaged assays.

A second apparent discrepancy concerns the effect of Nsp1 on translation initiation. Structural and biochemical studies show that the Nsp1 C-terminal domain occupies the ribosomal mRNA entry channel and can inhibit translation initiation^5,8,10^. Our measurements, however, suggest that the effect of Nsp1 on translation initiation is mRNA dependent. Initiation increased on the Host Reporter mRNAs that remained after Nsp1 expression, whereas other reporters (Host Reporter Long and the endogenous MYH9) showed a reduced fraction of translating mRNAs (even though the number of ribosomes per translation site was unchanged). This pattern is consistent with Nsp1 fully suppressing translation on a subset of mRNAs, while those that remain translationally active continue to support near-control levels of ribosome loading. Several mechanisms could account for this heterogeneity. First, extensive Nsp1-mediated depletion of cellular mRNAs may increase the availability of ribosomes and translation factors, partially offsetting direct inhibition of initiation on surviving transcripts. Second, reduced initiation on longer reporters could arise indirectly from their greater susceptibility to elongation-associated ribosome quality control rather than from stronger direct inhibition of initiation by Nsp1. RQC engagement can impose strong translational silencing, both by limiting subsequent initiation and by promoting degradation of the affected mRNA^44^. Finally, the magnitude of Nsp1-mediated initiation inhibition may itself depend on transcript-specific features, including UTR and coding-region architecture. These possibilities are not mutually exclusive, but collectively emphasize an important feature of our measurements: Nsp1 does not uniformly reduce initiation across individual mRNAs.

These findings raise the question of how SARS-CoV-2 RNAs remain productive while encoding Nsp1 themselves. During infection, genomic and subgenomic viral RNAs differ substantially in coding length but share common viral UTR elements^16,17^. Such differences could alter their relative exposure to the two routes identified here. Long viral coding regions provide a larger window in which Nsp1-mediated elongation defects could emerge, whereas continued cap-dependent initiation should allow Nsp1 to act repeatedly during initiation. These predictions remain speculative because our experiments used synthetic reporters rather than authentic viral RNAs, but they suggest that coding-region architecture, in addition to viral UTR identity, may contribute to determining Nsp1 susceptibility.

Our viral UTR reporters further suggest that no single viral UTR element is sufficient to escape Nsp1. Reporters containing both SARS-CoV-2 UTRs showed substantially increased translation output in the absence of Nsp1 but remained susceptible to Nsp1-mediated mRNA loss. This agrees with previous work in vitro showing that reporters containing both SARS-CoV-2 UTRs can have high intrinsic translation while remaining inhibited by Nsp1 to a similar degree as host reporters^10^. However, this differs from studies in which the SARS-CoV-2 5′ leader, particularly stem-loop 1, was sufficient to promote escape from Nsp1-mediated translational repression or cleavage^15,16^. One possible explanation is that Nsp1 sensitivity depends on the broader transcript context. In our reporters, altering the downstream coding region substantially changed Nsp1-mediated mRNA loss while leaving the upstream regulatory architecture unchanged. Effects attributed to the viral leader may therefore depend on interactions with the coding sequence, 3′ UTR, or other features of the experimental system. Authentic viral escape may similarly emerge from the complete viral RNA architecture and infection environment rather than from a single protective element.

Several limitations should be considered when interpreting our findings. First, single-molecule imaging provides direct measurements of mRNA abundance and translation dynamics but necessarily limits the number and diversity of transcripts that can be examined. Whether the mechanisms observed here extend broadly across the host transcriptome therefore remains unclear. Second, our measurements report the disappearance of intact mRNAs rather than the molecular events that initiate their degradation. We therefore cannot conclude that the initiation-associated pathway reflects direct Nsp1-mediated cleavage, even though biochemical studies have mapped Nsp1-dependent cleavage near the 5′ ends of mRNAs^11,12^. Likewise, ZNF598 dependence implicates ribosome-associated quality control but does not identify the underlying ribosomal substrate or downstream degradation machinery. Finally, ectopic Nsp1 expression in U2OS and HeLa cells may not fully reproduce the spatial, temporal, and molecular environment of authentic SARS-CoV-2 infection^42^.

Together, our results provide a framework for reconciling several previously distinct descriptions of Nsp1 activity. Rather than acting exclusively at translation initiation or during elongation, Nsp1 appears to create multiple opportunities for mRNA loss across the translation cycle. Cap-dependent initiation establishes an early route to mRNA depletion but also appears to create the conditions necessary for a later, ZNF598-sensitive pathway during elongation. This unexpected connection raises the possibility that Nsp1 perturbs ribosomes during cap-dependent initiation in a manner that influences their behavior long after they leave the start codon. How this state is established, how it alters ribosome dynamics during elongation, and how viral RNAs could tolerate these pathways remain important questions. Direct mapping of Nsp1-induced cleavage sites, transcriptome-wide measurements in ZNF598-deficient cells, and analysis of genomic and individual subgenomic RNAs during infection will be important for determining how broadly these mechanisms operate and how SARS-CoV-2 selectively maintains viral gene expression in the presence of Nsp1.

## METHODS

### Cell lines and cell culture

Human U2OS osteosarcoma cells, and Knockout ZNF598 U2OS were used for all single-mRNA reporter imaging experiments aside from those performed in HeLa cells with tagged MYH9 U2OS cells were maintained in Dulbecco’s modified Eagle’s medium (DMEM) supplemented with 10% fetal bovine serum, 1 mM L-glutamine, and 1% penicillin-streptomycin at 37 °C in a humidified 5% CO2 incubator. Cell density was maintained below confluence, and cells were routinely confirmed to be free of mycoplasma contamination. For live-cell NCT experiments, cells were plated on 35-mm glass-bottom dishes 1 day before imaging, typically at approximately 60-80% confluence.

For endogenous MYH9 experiments, an endogenously tagged MYH9 HeLa-Kyoto cell line previously generated and characterized by Wiggan et al.^39^ was used. This cell line contains biallelic C-terminal MYH9-mClover3-24×MS2 integrations and a monoallelic N-terminal 6×FLAG-mCherry2 insertion. Cells were maintained in high-glucose DMEM supplemented with 10% fetal bovine serum at 37 °C in a humidified 5% CO2 incubator. For experiments in this study, the tagged MYH9 cell line was transiently transfected with Nsp1, Nsp1ΔC, or control expression plasmids before imaging endogenous MYH9 mRNA and translation.

### Reporter design

Reporter plasmids were generated from established Nascent Chain Tracking (NCT) reporter architectures (Addgene #81085). In these reporters, the open reading frame (ORF) contains an N-terminal multi-epitope tag for nascent-chain detection (spaghetti monster HA)^45^, while the 3′ UTR contains 24×MS2 stem loops^21^ for visualization of individual reporter mRNAs. This design enables simultaneous imaging of nascent peptide synthesis and single reporter mRNAs in living cells ^19,20,31,33^. Reporter-specific changes to untranslated regions and coding sequences are described in the corresponding Results sections and figure schematics. Reporter variants were generated from the corresponding parental NCT reporter plasmid by restriction digestion at sites within the multiple cloning site to remove or linearize the sequence to be modified. The resulting plasmid backbone was assembled with the desired replacement sequence using Gibson Assembly and sequence confirmed with whole plasmid sequencing.

Reporter mRNAs and nascent chains were visualized using the All Probes Plasmid (APP) system described by Galindo et al.46. In this system, a single APP encodes the fluorescent probe components required for both mRNA and nascent-chain detection at defined relative expression levels. Reporter-based NCT experiments therefore used transient cotransfection of the reporter plasmid, the appropriate APP, and the indicated Nsp1 delivery construct. For the harringtonine experiments shown in Fig. 2F,G and Fig. S7E, cells were transfected with the CAAX-containing anti-HA StayGold/HaloTag APP (Addgene #241145). All other reporter-based imaging experiments used the corresponding untethered anti-HA StayGold/HaloTag APP described by Galindo et al.^46^ (APP 4), which lacks the CAAX domain.

Nsp1 delivery plasmids were generated by inserting the SARS-CoV-2 WT Nsp1 coding sequence into the common mammalian expression backbone used for the NCT reporter plasmids. To avoid directly tagging Nsp1, a porcine teschovirus 2A (P2A) sequence^25^ was placed downstream of Nsp1, followed by a fluorescent protein expressed as a separate translation product. For reporter-based NCT experiments, mCherry2 was used as the P2A-linked fluorescent marker to identify cells expressing the Nsp1 delivery construct. For endogenous MYH9 experiments, the same delivery strategy was used with eGFP as the downstream fluorescent marker. Where indicated, the corresponding Nsp1ΔC construct was generated and delivered using the same P2A-linked fluorescent-marker strategy.

### Transient transfection

All reporter-based imaging experiments were performed following transient cotransfection of the reporter construct, the corresponding All Probes Plasmid (APP), and either the Nsp1 or Nsp1ΔC expression plasmid. For most experiments, cells were transfected with the anti-HA/MCP-Halo APP derivative lacking the CAAX domain described in the **Reporter design** section. For the harringtonine experiments shown in **Fig. 2F,G** and **Fig. S7E**, the corresponding CAAX-containing APP (Addgene #241145; Galindo et al.^46^) was used.

Plasmids were transiently transfected using Plus Reagent and Lipofectamine LTX according to the manufacturer’s instructions. For each 35-mm imaging dish, the reporter, APP, and Nsp1 delivery plasmids were combined in a single transfection mixture. For all reporters except those containing the KDM5B open reading frame, 750 ng reporter plasmid was used, whereas 1,000 ng reporter plasmid was used for reporters containing the KDM5B open reading frame. In all experiments, 150 ng of APP plasmid and 200 ng of the corresponding Nsp1 delivery construct were included. The total DNA amount was 1.10 μg for fLuc-containing reporters and 1.35 μg for KDM5B-containing reporters. The combined DNA was diluted in Opti-MEM, and the amounts of Plus Reagent and Lipofectamine LTX were scaled to the total DNA amount according to the manufacturer’s recommendations. Transfection mixtures were incubated for 5–10 min at room temperature before addition to cells at approximately 50–70% confluence in 2 mL DMEM. Following addition of the transfection mixture, cells were incubated in DMEM containing the transfection reagents for approximately 12 h. Cells were then rinsed three times with unsupplemented phenol red-free DMEM and stained with 200 nM JFX650-HaloTag ligand in phenol red-free DMEM for 30 min at 37°C. After staining, cells were washed three additional times with unsupplemented phenol red-free DMEM to remove excess dye. The medium was then replaced with 2 mL supplemented phenol red-free DMEM for live-cell imaging. Cells were imaged 12–14 h after transfection, with imaging targeted to approximately 12 h after transfection.

### Live-cell single-molecule imaging of reporter translation

Live-cell NCT imaging of reporter constructs in U2OS cells was performed on a custom-built widefield fluorescence microscope configured for highly inclined and laminated optical sheet (HILO) illumination^19,47^. Briefly, excitation was provided by 488-, 561-, and 637-nm solid-state lasers coupled to a 60×/1.49 NA oil-immersion objective. Emission was detected on an aligned iXon Ultra 888 EMCCD camera. Cells were maintained during imaging in an enclosed stage-top incubator at 37°C with 5% CO2, and focus was stabilized using a CRISP autofocus system. Image acquisition was controlled using Micro-Manager^48^.

For reporter imaging experiments other than harringtonine runoff experiments, two-channel movies were collected to visualize reporter mRNAs and nascent-chain signals. Each volume consisted of 13 z-planes acquired at 0.5-μm spacing to cover the cytoplasmic volume. Individual images were acquired with a 53.64-ms exposure time. Full two-channel z-stacks were collected once per second for 20 s.

For experiments expressing the Nsp1 delivery construct, a single three-color volumetric image was acquired before time-lapse imaging. This initial image consisted of 13 z-planes collected in each channel and was used to document the Nsp1-associated marker. Subsequent time-lapse imaging was then performed using the reporter mRNA and nascent-chain channels.

Harringtonine runoff experiments (**Fig. 2F,G** and **Fig. S7E**) used the same volumetric imaging configuration, with 3 z-planes acquired at 0.5-μm spacing and a 53.64-ms exposure time per image. However, full image sets were collected at 1-min intervals for 35 minutes total to follow loss of translation-site signal after inhibition of new initiation.

### Image processing and spot quantification

Image analysis was performed using a custom Python pipeline incorporating TrackPy^50^, xarray, and a custom python package available on Github called CropArray^49^. For steady-state experiments, diffraction-limited mRNA puncta were detected in z-projected images of the MS2/Halo channel using TrackPy with a 5-pixel feature diameter. Integrated-intensity thresholds (min_mass) were optimized and manually checked for each reporter family using min_mass values between 500-2500 and subsequently held constant across matched control, Nsp1, and Nsp1ΔC conditions. Detected mRNA coordinates were used to generate three-dimensional crop arrays containing all imaging channels and time points.

Within each crop, the z-plane used for quantification was selected as the plane with the highest mean mRNA fluorescence within a centered 3-pixel-radius disk. The corresponding z-plane was then used for all imaging channels. Fluorescence intensity was measured as the mean signal within the centered 3-pixel-radius disk after subtraction of the mean local background measured from a one-pixel-wide outer annulus. Central puncta were independently identified in the mRNA and nascent-chain channels. When multiple features were detected within a crop, the feature nearest the crop center was retained. Only crops containing a central mRNA signal were included in subsequent analyses, and an mRNA was classified as translating when a coincident central nascent-chain signal was detected. This coordinate-matched strategy limited the assignment of unrelated protein puncta as translation sites.

### Steady-state reporter analysis

Steady-state measurements were made from cells imaged 11–14 h after transfection, with imaging targeted to approximately 12 h after transfection. For each cell, the total number of mRNAs, the number of translation sites, and the fraction of translating mRNAs were calculated at each time point and averaged over the first five frames. The translating fraction was defined as the number of mRNAs associated with a detectable nascent-chain signal divided by the total number of detected mRNAs. Mean mRNA and translation-site intensities were calculated from signal-positive puncta over the same time window.

For normalized comparisons, individual cell measurements were divided by the mean of the matched No Nsp1 condition from the same biological replicate. Imaging dates were treated as independent biological replicates, and replicate-level means were used for superplot visualization and statistical analysis. Cell-level outliers were identified separately within each reporter and experimental condition using Tukey’s interquartile range criterion of 1.5 × IQR. Cell-level outlier identification and exclusion were performed as described in the **Statistical analysis** section.

### Translation inhibitor treatments

To confirm that detected nascent-chain puncta represented actively translating ribosomes, cells containing visible translation sites were imaged before and after puromycin treatment. Up to five cells were imaged simultaneously at 1-min intervals, with five pre-treatment time points collected to establish a baseline. Puromycin was then added to a final concentration of 50 μg/mL, and imaging continued at the same interval for up to 15 min. Translation-site fluorescence was quantified as described in the **Image processing and spot quantification** section. Translation-site signals were averaged within each cell at each time point, and traces were aligned to the time of puromycin addition.

For Harringtonine runoff experiments, cells containing visible translation sites were imaged at 1-min intervals for five pre-treatment time points before addition of harringtonine to a final concentration of 3 μg/mL. Imaging then continued at the same interval for 40 min after drug addition. Translation-site fluorescence was quantified as described in the **Image processing and spot quantification** section. Normalization, temporal alignment, and runoff fitting are described in the **Harringtonine runoff analysis** section.

### Harringtonine runoff analysis

For harringtonine experiments, translation sites were identified before drug addition, and centered three-dimensional crops containing all imaging channels and time points were generated using CropArray^49^. Translation-site fluorescence was measured throughout each movie, summed for each cell, and normalized to the mean fluorescence signal from the final three pre-treatment frames.

Normalized runoff trajectories were fit by nonlinear least squares using a hyperbolic tangent model modified to include a nonzero residual fluorescence fraction *r*:

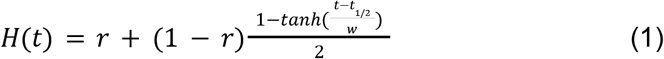

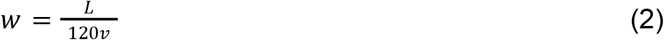

where *H*(*t*) is normalized translation-site fluorescence, *t* is the residual fluorescence fraction, 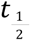 is the fitted midpoint of the runoff transition, *w* is the fitted transition width, *L* is the coding-region length in amino acids (780 aa), *v* is the elongation rate in amino acids s⁻¹. Harringtonine was added after the 5-min pre-treatment baseline, and the first minute after drug addition was excluded to allow inhibition of new initiation. To account for the delay between harringtonine addition and effective inhibition of initiation, post-treatment time was offset by 1 min. The first image acquired 1 min after drug addition was therefore treated as the start of the run-off period for fitting.

Ribosome initiation rates were calculated for individual cells from the calibrated ribosome occupancy and fitted elongation rate:

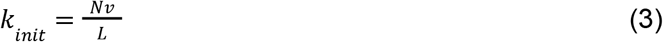

where *N* is the estimated ribosome occupancy per translation site, and *k_init_* is the ribosome initiation rate.

Harringtonine run-off traces were normalized to the mean signal from the final three pre-treatment frames and fit to the modified hyperbolic tangent model described above. The first minute after harringtonine addition was excluded to account for the delay before inhibition of new initiation. Harringtonine-resistant translation sites were analyzed separately from the run-off fits and were defined as sites retaining at least 5% of their mean pre-treatment signal at the endpoint.

The harringtonine-resistant fraction was measured independently from the runoff fit. Translation sites present during the five pre-treatment frames were compared with sites remaining at the designated endpoint. Sites retaining at least 5% of the mean pre-treatment translation-site intensity were classified as harringtonine-resistant, and the harringtonine-resistant fraction was calculated as the number of retained sites divided by the mean number of translation sites present during the pre-treatment baseline.

### Nascent-chain calibration and ribosome occupancy estimation

Translation-site fluorescence was converted to estimated ribosome occupancy using the previously calibrated KDM5B reporter. Koch et al.^33^ estimated an average occupancy of 14.6 ± 5.6 ribosomes per actively translating KDM5B mRNA. KDM5B and Host Reporter control cells were imaged on the same day using the acquisition settings described in the **Live-cell single-molecule imaging of reporter translation** section. For each reporter, translation-site intensities were averaged within each biological replicate. These replicate means were then averaged across replicates, so that each biological replicate contributed equally, regardless of the number of cells analyzed.

The mean KDM5B translation-site intensity calculated from this replicate-averaged calibration, *I_KDM_*_5*B*_, was assigned an occupancy of 14.6 ribosomes. The Host Reporter measured in the same calibration experiment was then used to transfer the KDM5B-derived absolute scale to the Host Reporter. Its ribosome occupancy was calculated as follows:

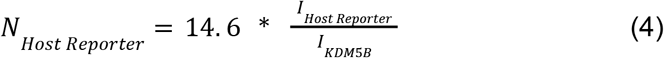

where *N_Host Reporter_* is the estimated ribosome occupancy per actively translating Host Reporter mRNA, *I_Host Reporter_* is the replicate-averaged mean Host Reporter translation-site intensity, and *I_KDM5B_* is the replicate-averaged mean KDM5B translation-site intensity measured under matched imaging conditions.

### Local mRNA pair-enrichment analysis

For **Fig. 1E**, local mRNA organization was quantified from cytoplasmic mRNA positions during the first five imaging frames. Frames containing fewer than 10 cytoplasmic mRNAs were excluded, and cells were required to contain at least three valid frames. Cumulative numbers of mRNA pairs were measured over distances from 0.78 to 1.56 μm in 0.13-μm increments and compared with 199 Monte Carlo simulations. Simulated distributions preserved cell shape, nuclear exclusion, total mRNA number, the number of mRNAs within each radial region, and a minimum mRNA-mRNA separation of 0.65 μm corresponding to the 5-pixel detection diameter. Local pair enrichment was calculated as the ratio of the area under the observed pair-count curve to the mean area under the simulated pair-count curves over the analyzed distance range. An enrichment value of 1 therefore represents the matched random expectation. Frame-level enrichment values were averaged within each cell and then within each biological replicate.

### mRNA tracking and mean-squared displacement analysis

For mRNA mobility measurements in **Fig. 1C**, cytoplasmic mRNA detections from retained cells were linked between frames using TrackPy^50^. Linking was performed in the xy plane with a maximum displacement of 7 pixels between consecutive frames and a memory of one missing frame; z positions were retained for subsequent analysis but were not used for linking. Only trajectories spanning at least five frames were analyzed. Coordinates were converted to physical units using an xy pixel size of 0.130 μm and z-step size of 0.500 μm, and lag times were calculated using the acquisition interval for each condition. Time-averaged mean-squared displacement (MSD) was calculated from displacement pairs separated by the specified lag, with missing frames omitted. Lags of up to 10 frames were analyzed, with at least three displacement pairs required per trajectory. Track-level MSD values were averaged equally within each cell, followed by equal averaging of cells within each biological replicate. Group MSD curves represent the mean ± standard error of the mean (SEM) across biological replicates.

### Labeling of endogenous MYH9 mRNA and translation sites

For imaging of the endogenously tagged MYH9 line, endogenous MYH9 mRNAs were visualized through HaloTag-MCP binding to the 24×MS2 cassette. As described by Wiggan et al.^39^, cells with relatively low HaloTag-MCP expression were incubated with 200 nM JF646 HaloTag ligand for 20 min before imaging. Endogenous MYH9 translation sites were detected using anti-FLAG immunofluorescence in fixed-cell experiments. Because mature mCherry2 and mClover3 fluorescence require chromophore maturation, nascent translation sites were identified by colocalization of the anti-FLAG Fab signal and the reporter mRNA signal, in the absence of mature fluorescent protein signal (mClover3 and mCherry2).

### Spinning-disk confocal imaging of endogenously tagged MYH9 cells

Imaging of the endogenously tagged MYH9 line was performed using the confocal workflow described by Wiggan et al^39^. Images were acquired on an Olympus IX8 spinning-disk microscope equipped with a CSU22 head, 405-, 488-, 561-, and 640-nm laser lines, and an iXon Ultra 888 EMCCD camera controlled with SlideBook software. A 100×/1.40 NA or 60×/1.42 NA objective was used depending on the experiment. Cells were imaged in glass-bottom dishes in an environmental enclosure maintained at 37°C with 5% CO2.

### Endogenous MYH9 image analysis

For analysis of the endogenously tagged MYH9 cell line, image processing and quantification followed the pipeline described by Wiggan et al. Cell, nuclear, and cytoplasmic segmentations were generated using Cellpose^51^. Translation sites were identified from maximum-intensity projections of the anti-FLAG channel using Ilastik-based pixel classification^52^, followed by segmentation in ImageJ to obtain centroid coordinates. Translation-site intensities were then measured from a central circular region of interest with local background subtraction. MYH9 mRNAs were detected and quantified in 3D from the MS2/Halo channel using FISH-quant v2 following Laplacian of Gaussian filtering and spot detection^53^. Translation sites and mRNAs were then assigned to individual cells and nuclei using the corresponding Cellpose segmentations. All endogenous MYH9 measurements reported in this study were derived from this analysis framework.

### Statistical analysis

Statistical analyses were performed in Python using pandas, NumPy, SciPy, and statsmodels. Unless otherwise stated, data are reported as mean ± SEM across independent biological replicates, defined by imaging date. Cell-level measurements were averaged within each biological replicate, and replicate means were used for statistical testing and visualization. Individual cells are shown to illustrate population distributions but were not treated as independent biological replicates.

For normalized steady-state measurements, individual cell values were normalized to the mean of the matched No Nsp1 condition from the same biological replicate before replicate-level averaging. Normalized conditions were compared with the reference value of 1 using two-sided one-sample t-tests, whereas matched experimental conditions were compared using two-sided paired t-tests on replicate means. Cell-level outliers were identified separately within each reporter and condition using Tukey’s 1.5 × interquartile range criterion. Within each biological replicate, the distributions of cell-level mRNA counts and translation-site counts were evaluated using Tukey’s 1.5 × IQR criterion. Cells falling outside the resulting range for either metric were classified as outliers and excluded at the cell level, with all measurements from those cells removed from subsequent analyses.

For local mRNA pair-enrichment analysis, frame-level values were averaged within cells and then across biological replicates, with an enrichment value of 1 representing the matched spatial-randomization expectation. For MSD analysis, track-level values were averaged within cells and then within biological replicates, and group curves are shown as mean ± SEM across replicates. Statistical significance was defined as p < 0.05; exact p values are shown where appropriate, and ns indicates p > 0.05.

## ACKNOWLEDGEMENTS

We thank the Luke Lavis lab for the HaloTag ligands used in this work. We thank the Simon Bekker-Jensen lab for the knockout ZNF598 U2OS cell line. We thank members of the Stasevich lab for helpful discussions. We used large language models (Claude, Anthropic; ChatGPT, OpenAI) to improve the clarity and grammar of the manuscript. All text was reviewed and edited by us, and we take full responsibility for the contents.

## AUTHOR CONTRIBUTIONS

Conceptualization: A.K, and T.J.S. Methodology: A.K, T.J.S, and O.W. Software: A.K, and T.J.S. Validation: A.K. Formal analysis: A.K. Investigation: A.K, E.B, and O.W. Resources: T.J.S. Data curation: A.K and T.J.S. Writing drafts A.K and T.J.S: Review and editing: A.K, E.B, O.W., C.D.S, T.M., B.J.G., C.D.S., and T.J.S. Visualization: A.K, E.B, O.W, C.D.S, and T.J.S. Supervision: T.J.S. Project administration: T.J.S. Funding acquisition: C.D.S., B.J.G.,and T.J.S.

## DATA AND MATERIAL AVAILABILITY

Code for generating will be made available on GitHub and FigShare upon publication.

