## Supplemental Figures for "Live-cell, single-molecule imaging resolves the dual nature of mRNA silencing by SARS-CoV-2 Nsp1"

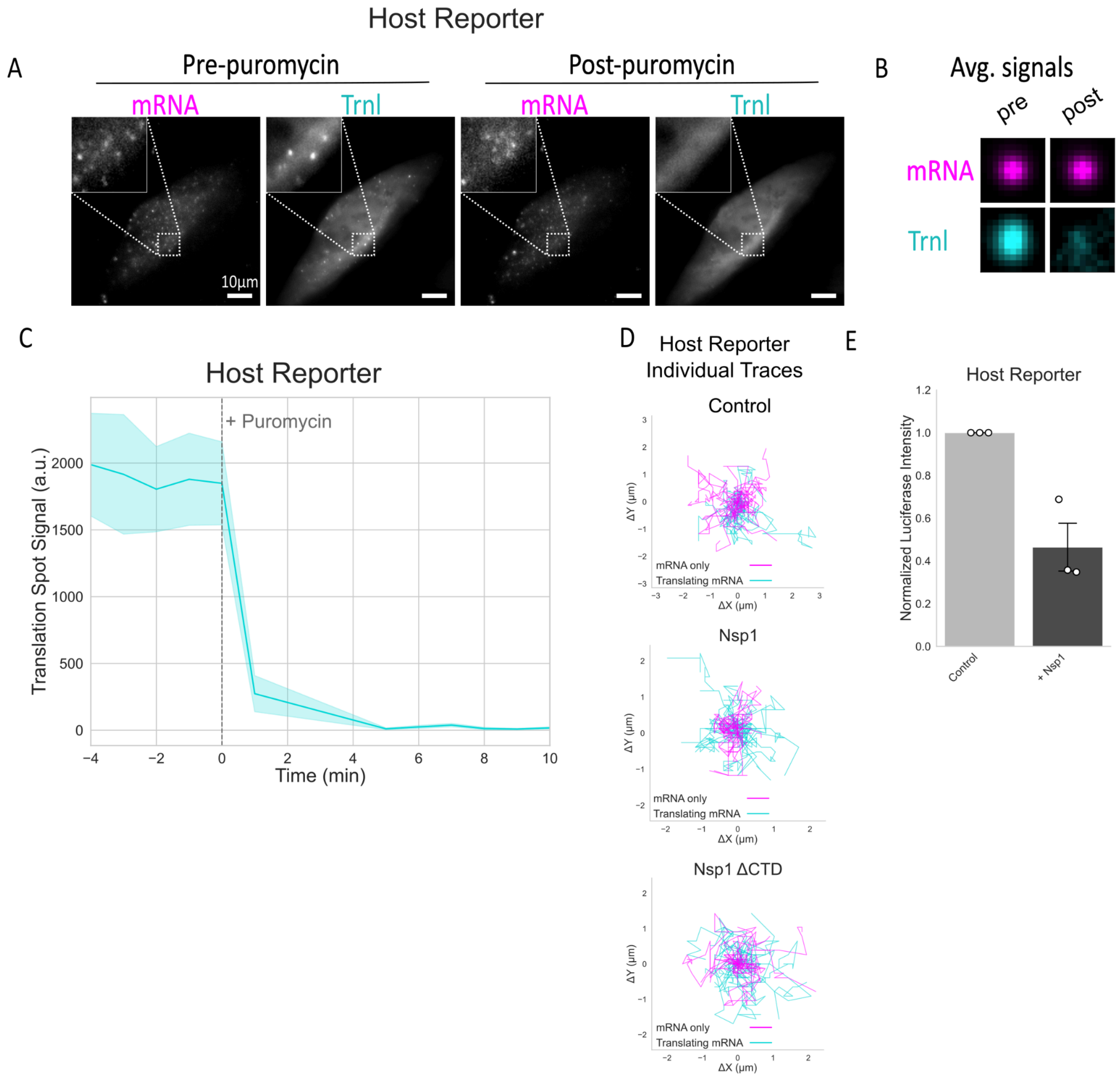


**Figure S1. Validation of nascent-chain translation in the Host Reporter.**
(A) Representative images of reporter mRNA (magenta) and nascent translation signal (cyan) before and after puromycin addition. The insets show zoomed-in views of reporter mRNA and nascent-chain signals. (B) Average mRNA and translation-site signals before and after puromycin treatment. (C) Mean translation-site fluorescence over time relative to puromycin addition. The solid line indicates the mean and shading indicates the standard error of the mean. (D) Random selection of 25 mRNA and translation traces plotted with start points at (0,0). Scale bars, 10 μm.

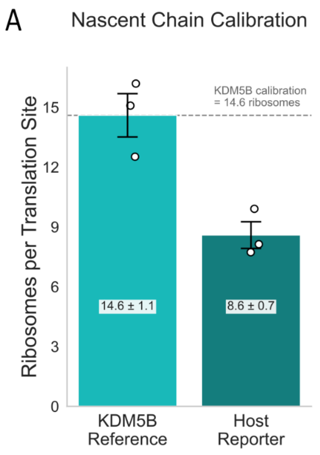


**Figure S2. Nascent-chain calibration of Host Reporter ribosome occupancy.**
(A) Nascent-chain signal from the KDM5B calibration reporter and Host Reporter used to estimate ribosome occupancy. Estimated occupancies were 14.6 ± 1.1 (mean ± SEM) ribosomes per translation site for the KDM5B reference and 8.6 ± 0.7 ribosomes per translation site for the Host Reporter. See Methods for details of this calibration.

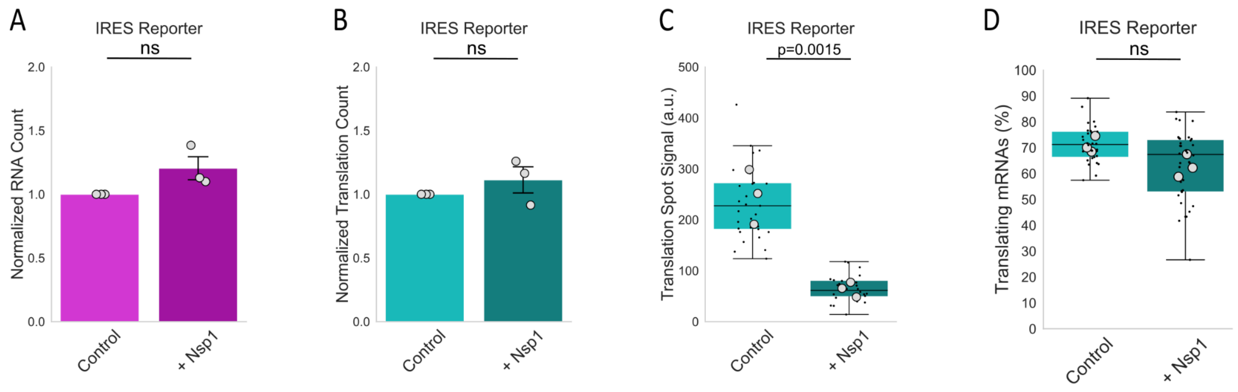


**Figure S3. Nsp1 alters translation signal brightness but not mRNA abundance of the IRES Reporter.**
(A) Normalized IRES Reporter mRNA abundance in control and Nsp1-expressing cells. (B) Normalized number of translation sites per cell. Translation-site number with Nsp1 was 111 ± 10.3% (mean ± SEM) of control. (C) Translation-site intensity of actively translating mRNAs. Translation-site intensity was 247 ± 31.1 a.u. in control cells and 63.7 ± 8.4 a.u. with Nsp1. (D) Fraction of reporter mRNAs undergoing translation in control and Nsp1-expressing cells. P values were calculated using two-sided one-sample t-test for normalized conditions, or paired two-sided t-test for matched conditions across biological replicates. ns indicates p>0.05. Panels A and D are reproduced from the main text for side by side comparison.

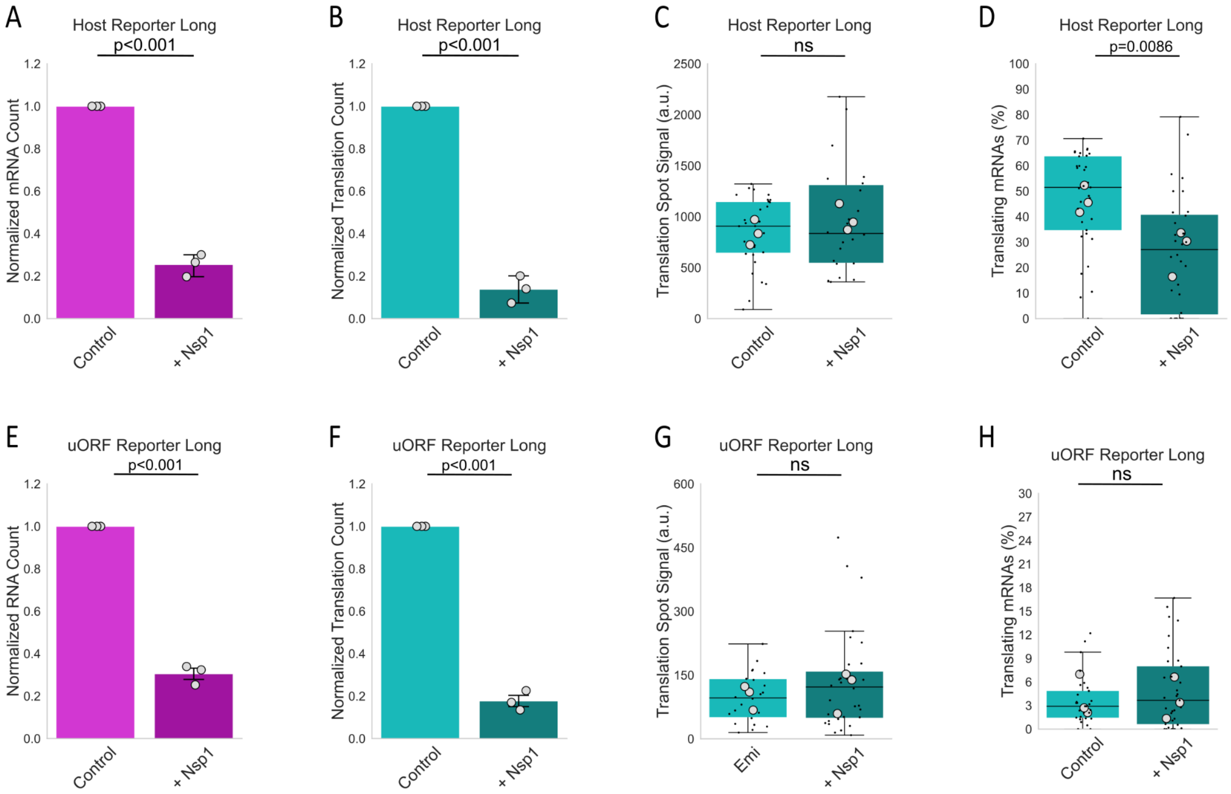


**Figure S4. Nsp1 strongly suppresses long KDM5B-containing reporters.**
(A–D) Host Reporter Long measurements in control and Nsp1-expressing cells. (A) Normalized mRNA abundance. (B) Normalized translation-site number. Translation-site number with Nsp1 was 13.8 ± 3.7% (mean ± SEM) of control. (C) Translation-site intensity. Translation-site intensity was 843 ± 71.5 a.u. in control cells and 982 ± 76.2 a.u. in Nsp1-expressing cells. (D) Translating fraction in control and Nsp1-expressing cells. (E–H) Corresponding measurements for uORF Reporter Long. (E) Normalized mRNA abundance. (F) Normalized translation-site number. Translation-site number with Nsp1 was 17.7 ± 2.7% of control. (G) Translation-site intensity. Translation-site intensity was 100 ± 16.7 a.u. in control cells and 117 ± 29 a.u. with Nsp1. (H) Translating fraction in control and Nsp1-expressing cells. P values were calculated using two-sided one-sample t-test for normalized conditions, or paired two-sided t-test for matched conditions across biological replicates. ns indicates p>0.05. Panels A, D, E, and H are reproduced from the main figure for side by side comparison.

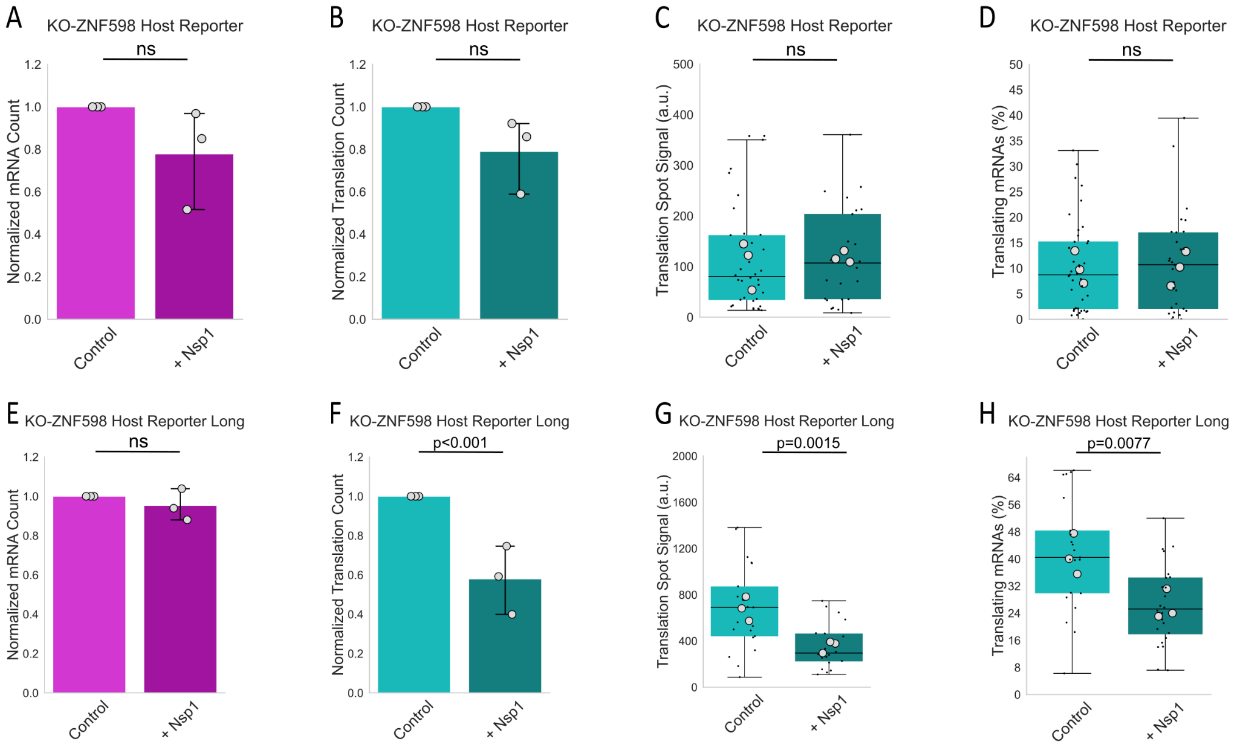


**Figure S5. Nsp1 responses of short and long reporters in ZNF598-knockout cells.**
(A–D) Host Reporter measurements in ZNF598-knockout U2OS cells. (A) Normalized mRNA abundance in control and Nsp1-expressing cells. (B) Normalized translation-site number. Translation-site number with Nsp1 was 79 ± 10.2% (mean ± SEM) of control. (C) Translation-site intensity. Translation-site intensity was 109 ± 27.4 a.u. in control cells and 119 ± 6.66 a.u. with Nsp1. (D) Translating fraction in control and Nsp1-expressing cells. (E–H) Corresponding measurements for Host Reporter Long. (E) Normalized mRNA abundance in control and Nsp1-expressing cells. (F) Normalized translation-site number. Translation-site number with Nsp1 was 57.9 ± 10.0% of control. (G) Translation-site intensity. Translation-site intensity was 680 ± 60.5 a.u. in control cells and 355 ± 30.1 a.u. with Nsp1. (H) Translating fraction in control and Nsp1-expressing cells. P values were calculated using two-sided one-sample t-test for normalized conditions, or paired two-sided t-test for matched conditions across biological replicates. ns indicates p>0.05. Panels A, D, E, and H are reproduced from the main figure for side by side comparison.

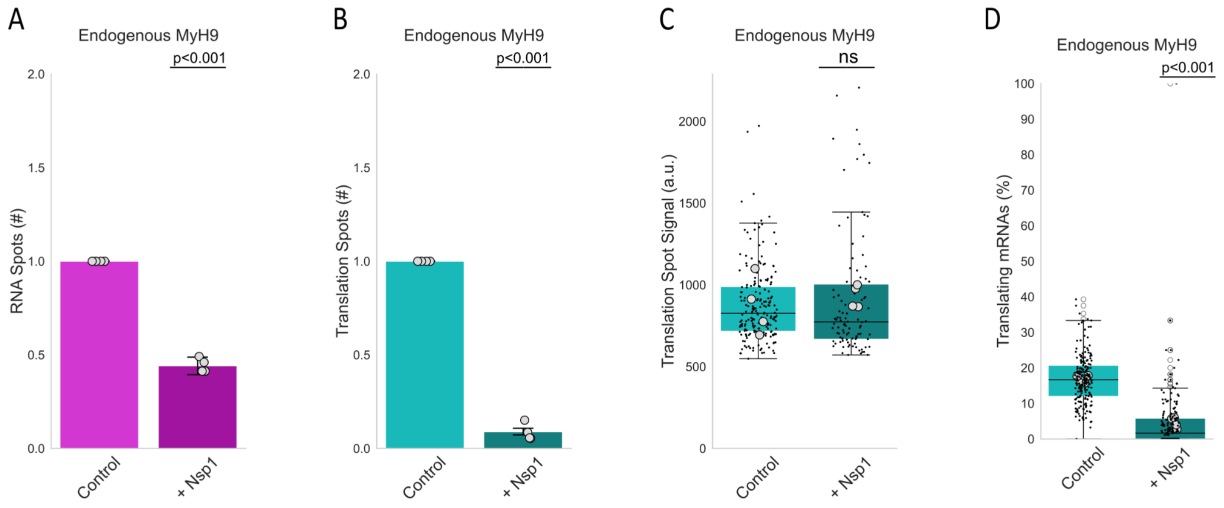


**Figure S6. Nsp1 impacts Endogenous MYH9 system similar to Host Reporter Long.**
(A) Normalized endogenous MYH9 mRNA abundance in control and Nsp1-expressing cells. (B) Normalized number of translation sites per cell in control and Nsp1-expressing cells. (C) Translation-site intensity of actively translating mRNAs. Translation-site intensity was 871 ± 88.8 a.u. (mean ± SEM) in control cells and 929 ± 35.1 a.u. with Nsp1. (D) Fraction of endogenous MYH9 mRNAs undergoing translation in control and Nsp1-expressing cells. P values were calculated using two-sided one-sample t-test for normalized conditions, or paired two-sided t-test for matched conditions across biological replicates. ns indicates p>0.05. Panels A, B, and D are reproduced from the main text for side by side comparison.

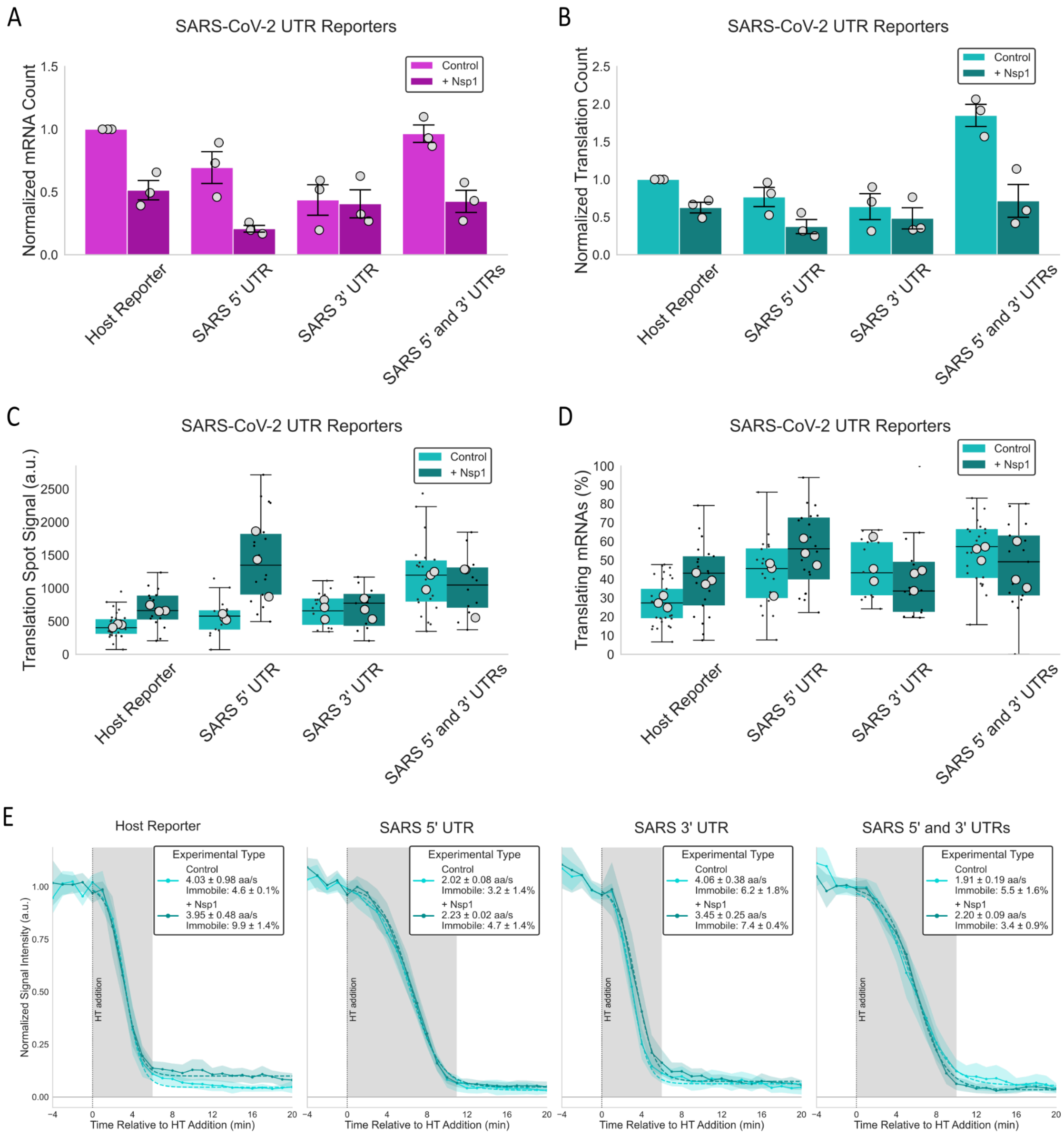


**Figure S7. SARS-CoV-2 UTRs alter reporter translation dynamics.**
(A–D) Steady-state measurements of the Host Reporter and reporters containing the SARS-CoV-2 5′ UTR, 3′ UTR, or both viral UTRs in control and Nsp1-expressing cells. (A) Normalized mRNA abundance. (B) Normalized translation-site number. (C) Translation-site intensity. Translation-site intensity for the combined UTR reporter was 1140 ± 83.3 a.u. (mean ± SEM) compared with 437 ± 15.7 a.u. for the Host Reporter. (D) Translating fraction for each reporter in control and Nsp1-expressing cells. (E) Mean normalized translation-site intensity following harringtonine addition for each reporter. Elongation rates were: SARS 5′ UTR, 2.0 ± 0.1 aa $s^{-1}$; + Nsp1, 2.23 ± 0.02 aa $s^{-1}$; SARS 3′ UTR, 4.1 ± 0.4 aa $s^{-1}$; + Nsp1, 3.5 ± 0.3 aa $s^{-1}$; SARS 5′ & 3′ UTR, 1.9 ± 0.2 aa $s^{-1}$; + Nsp1, 2.2 ± 0.1 aa $s^{-1}$. HT-resistant fractions are reported within the plots. Solid lines indicate mean runoff profiles, colored shading indicates variability, and the shaded gray region approximates runoff times. Panels A and B are reproduced from the main text for side by side comparison.
